# Theory of site-specific DNA-protein interactions in the presence of nucleosome roadblocks

**DOI:** 10.1101/214387

**Authors:** R. Murugan

## Abstract

We show that nucleosomes can efficiently control the relative search times spent by transcription factors (TFs) on one- (1D) and three-dimensional (3D) diffusion routes towards locating their cognate sites on DNA. Our theoretical results suggest that the roadblock effects of nucleosomes are dependent on the relative position on DNA with respect to TFs and their cognate sites. Especially, nucleosomes exert maximum amount of hindrance to the 1D diffusion dynamics of TFs when they are positioned in between TFs and their cognate sites. The effective 1D diffusion coefficient (*χ*_*TF*_) associated with the dynamics of TFs in the presence of nucleosome decreases with the free energy barrier (*µ*) associated the sliding dynamics of nucleosomes as 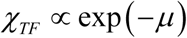. Subsequently the mean first passage time (*η*_*L*_) that is required by TFs to scan *L* number of binding sites on DNA via 1D diffusion increases with *μ* as 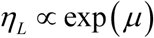. When TFs move close to nucleosomes then they exhibit a typical sub-diffusive dynamics. Nucleosomes can enhance the search dynamics of TFs when TFs present in between nucleosomes and transcription factor binding sites (TFBS). The level of enhancement effects of nucleosomes seems to be much lesser than the level of retardation effects when nucleosomes present in between TFs and their cognate sites. These results suggest that nucleosome depleted regions around the cognate sites of TFs is mandatory for an efficient site-specific interactions of TFs with DNA. Remarkably the genome wide positioning pattern of TFs shows maximum at their specific binding sites and the positioning pattern of nucleosome shows minimum at the specific binding sites of TFs under *in vivo* conditions. This seems to be a consequence of increasing level of breathing dynamics of nucleosome cores and decreasing levels of fluctuations in the DNA binding domains of TFs as they move across TFBS. Since the extent of breathing dynamics of nucleosomes and fluctuations in the DBDs of TFs are directly linked with their respective 1D diffusion coefficients, the dynamics of TFs becomes slow as they approach their cognate sites so that TFs form tight site-specific complex. Whereas the dynamics of nucleosomes becomes rapid so that they pass through the cognate sites of TFs. Several *in vivo* datasets on genome wide positioning pattern of nucleosomes as well as TFs seem to agree well with our arguments. We further show that the condensed conformational state of DNA can significantly decrease the retarding effects of nucleosome roadblocks. The retarding effects of nucleosomes on the 1D diffusion dynamics of TFs can be nullified when the degree of condensation of the genomic DNA is such that it can permit a jump size associated with the dynamics of TFs beyond *k* > 150 bps.

## 1. Introduction

Finding a specific site on the genomic DNA in the presence of enormous amount of nonspecific binding sites is one of the fundamental random search problems in biological physics. Transcription factors (**TFs**) precisely regulate the cellular expression levels of several genes across prokaryotes and eukaryotes by site specifically binding with the corresponding *cis*-regulatory modules (**CRMs**) present on the genomic DNA [1-4]. Site-specific binding of TFs with DNA was initially thought as a single step three-dimensional (**3D**) diffusion controlled collision process. Several *in vitro* studies on the binding kinetics of *lac*-repressor with its Operator DNA sequence revealed a bimolecular site-specific collision rate in the order of ~10^9^-10^10^ M^−1^s^−1^ which is ~10-10^2^ times faster than the 3D diffusion controlled collision rate limit. This rapid searching phenomenon was successfully explained by Berg et.al., [5, 6] using a two-step mechanism in which the TF molecule first binds with DNA in a nonspecific manner via 3D diffusion route and then searches for its cognate site via various one-dimensional (**1D**) facilitating mechanisms such as sliding, hopping and intersegmental transfers along DNA. In sliding, TFs moves on DNA with unit base-pair step size. We define 1 base-pair (bps) ~ 3.4 x 10^−10^ m and we measure the length variables in terms of bps throughout this paper. TFs move along the DNA polymer with few base-pairs step size in the hopping mode of dynamics and few hundred to few thousand base-pairs step-size during intersegmental transfer dynamics (**Figs. 1A** and **B**). Intersegmental transfers occur when two distal segments of the same DNA polymer come close over 3D space via ring closure events [7-9].

**FIG. 1.**
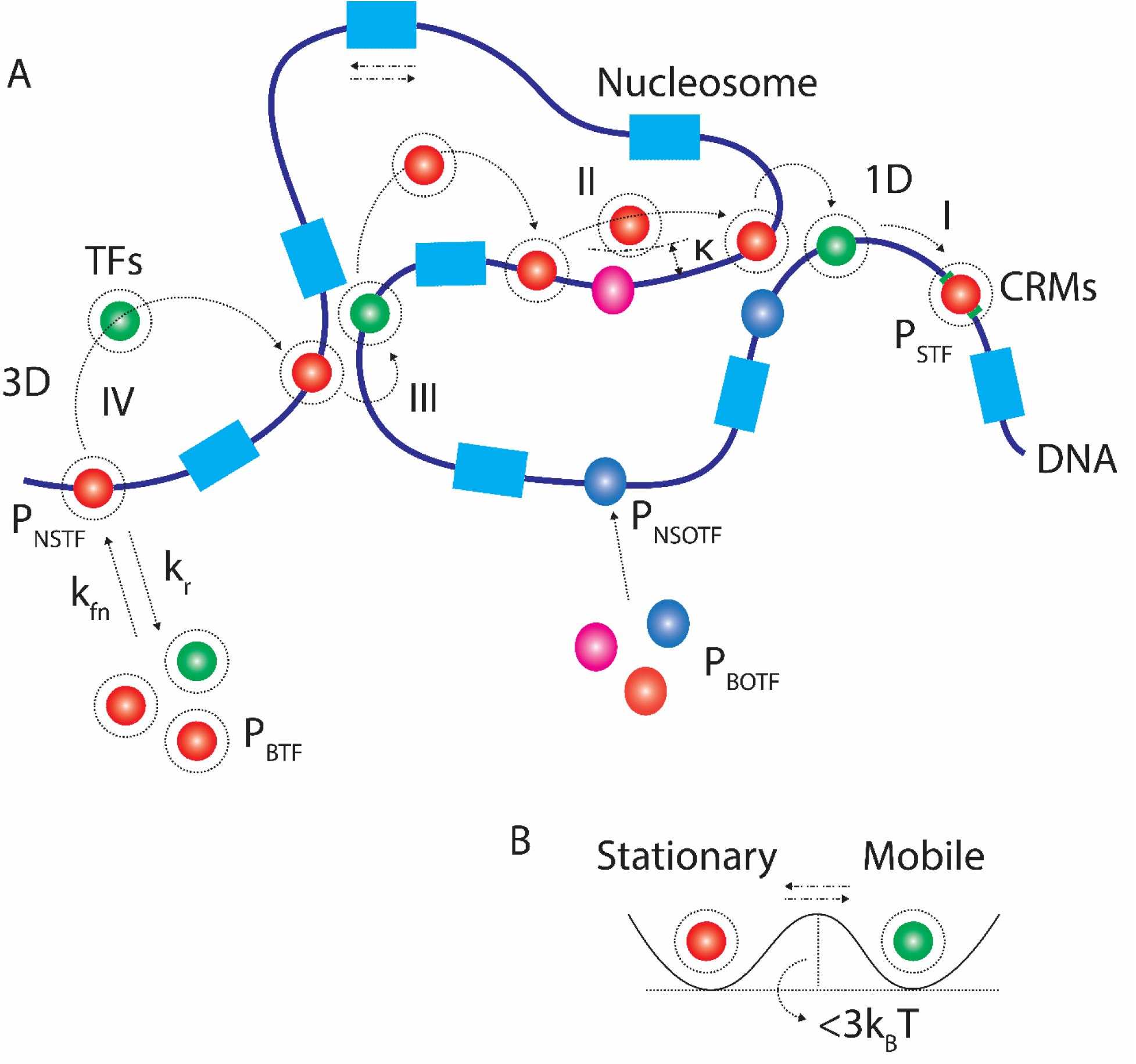
**A**. Various factors influencing the random search dynamics of TFs (*P*_*BTF*_) towards their CRMs on the genomic DNA. In a standard two step model, the TF of interest first non-specifically binds with DNA with a rate constant *k*_*fn*_ and then searches for the specific binding site via 1D diffusion along the DNA lattice and subsequently form the site specific complex (*P*_*STF*_). Before reaching CRMs, TFs undergo several cycles of associations and dissociations where *k*_*r*_ is the dissociation constant. Here the 1D diffusion of TFs will be facilitated by sliding (**I**), hopping (**II**) and intersegmental transfers (**III**). The TF molecule moves with unit base-pair step size in sliding and few bps in hopping. When two distal segments of the same DNA polymer come close over 3D space via ring closure events, intersegmental transfers occur (**III**). All these facilitating processes are defined well within the framework of 1D diffusion since they are confined within the Onsager radius (*κ*) during these processes. Here Onsager radius is defined as the distance between the charged molecules at which the overall electrostatic interaction energy will be comparable with that of the background thermal energy. Here the nonspecific binding is mainly driven by the electrostatic attractive forces operating at the DNA-protein interface where the backbone of DNA is negatively charged due to the presence of phosphate groups and the DNA binding domains of TFs is positively charged due to the presence of basic amino acid side chains. The 1D diffusion dynamics of TFs of interest along the DNA lattice under *in vivo* conditions will be impeded by the presence of dynamic roadblocks (*P*_*BOTF*_) such as other TFs and semi stationary roadblocks such as nucleosomes. Here nucleosomes are semi stationary massive roadblocks which diffuse or translocate much slower than TFs. **B**. Apart from these factors the DNA binding domains of TFs undergo fluctuations between stationary and mobile states where the free energy barrier associated with these fluctuations seems to be < 3*k*_*B*_*T*.

Site specific binding of TFs with the genomic DNA seems to be influenced by several factors under *in vivo* conditions [9] viz. (a) conformational state of DNA [9, 10] (b) spatial organization of various functionally related CRMs along the genomic DNA [11, 12], (c) presence of other dynamic roadblock TF proteins and semi-stationary roadblocks such as nucleosomes especially in eukaryotes [13-16], (d) naturally occurring sequence traps on DNA [17, 18], (e) conformational fluctuations in the DNA binding domains (**DBDs**) of TFs (**Fig. 1B**) [19-21] and (f) the nonspecific electrostatic attractive forces and the counteracting shielding effects of other solvent ions and water molecules acting at the DNA-protein interface [22]. Several theoretical [8, 9, 17, 20, 23], computational [24-27] and experimental studies have been carried out to understand the effects of various factors **a-f** on the site specific binding kinetics of TFs with DNA.

In general, the efficiency of TFs in locating their cognate sites on DNA via random searching mechanism seems to be strongly dependent on the relative amount of times spent on 3D and 1D diffusion routes [11, 20]. Clearly neither pure 1D nor pure 3D diffusion is efficient mode of searching for the specific binding sites [9, 11]. Under ideal situation, maximum efficiency of random searching can be achieved only when TFs spend equal amount of times in both 1D as well as 3D diffusions [8, 11]. This trade off balance of 1D and 3D diffusion times will be modulated by the presence of factors **a-f**. For example presence of roadblocks or sequence traps warrants more dissociations and 3D excursions of TFs rather than 1D sliding along DNA. Relaxed conformational state of DNA enhances the sliding type dynamics of TFs rather than hopping and intersegmental transfers and so on. In eukaryotic systems nucleosomes play special roles in modulating the relative search times associated with the binding of TFs with their CRMs on the chromosomal DNA segments [13-16, 28, 29]. Nucleosomes are slowly diffusing massive roadblocks which are the main sources of epigenetic variations in eukaryotic systems [3].

Each nucleosome core particle dynamically spans around 147 bps of the genomic DNA which subsequently become transiently inaccessible for the nonspecific or specific binding of most of the TFs [3, 28]. The length of the linker DNA between two consecutive nucleosome particles seems to be in the range of ~10-100 bps [29, 30]. Nucleosomes affect the overall searching dynamics of TFs towards their CRMs at least in two different ways viz. they (a) dynamically control the accessibility of specific as well as non-specific binding sites and, sequence mediated traps towards incoming TFs from the bulk cytoplasm and (b) introduce semi-stationary roadblocks across the 1D diffusion dynamics of TFs along the DNA polymer [16, 31]. When CRMs are concealed by the bulky nucleosome complex, then TFs wander over nearby or some other location of the same DNA chain or wait until the slow dislocation of the nucleosome assembly [9]. Otherwise some other active or competitive passive machinery is required to dismantle the nucleosome complexes from the specific binding sites of TFs. Such active displacement or disassembling of nucleosomes always incurs energy input in the form of ATP hydrolysis. Particularly the chromatin remodeling factor such as ACF (ATP-dependent chromatin-remodeling factor) is required to increase the accessible DNA lengths [28, 32] for the binding of TFs in eukaryotes. Depending on the relative timescales associated with the 1D diffusion dynamics of TFs or kinetics of nucleosome dissociation and dislocation, the underlying random search process of TFs will be either diffusion-controlled, kinetic-controlled or a combination of them.

The overall time required by TFs to locate their cognate sites will be approximately equal to the sum of the random search time and the time required for the displacement or dissociation of nucleosomes from the cognate sites of TFs. This is similar to the coupled transcription-splicing [33] where the small nuclear ribonucleoproteins (**snRNPs**) which are performing 1D diffusion on the already emerged out segment of pre-mRNA need to wait until the appearance of splicing sites from the transcription assembly to initiate the splicing process. Most of the earlier works on the role of nucleosomes on gene regulation were mainly focused on (1) unravelling the sequence dependent statistical distribution of nucleosomes over several locations of the genomic DNA [13, 28, 30, 31, 34, 35] and prediction of such patterns by considering the data obtained from *in vivo* MNase-seq experiments [16, 31] and (2) understanding the mesoscopic mechanism underlying the sliding dynamics of nucleosomes in the presence and absence of other DNA binding ligands and associated potentials [36-39]. In this context there are several open questions viz. (**a**) how exactly the slowly diffusing massive roadblocks such as nucleosomes dynamically influence the overall search times required by TFs to locate their cognate sites on the genomic DNA, (**b**) how the conformational state of DNA influences the interplay between the dynamics of TFs and nucleosomes and (**c**) how the nucleosome positioning or occupancy pattern around the transcription state sites (**TSS**) or transcription factor binding sites (**TFBS**) on the genomic DNA influences the overall search times required by TFs to find their cognate sites. Particularly the dynamic interaction between the random search mechanism of TFs and the slow diffusion as well as breathing actions of nucleosomes are not clearly understood. In this paper we address these questions using a combination of theoretical and simulation tools in detail.

## 2. Theory

Let us consider a linear DNA lattice of size *N* bps containing the CRM corresponding to a TF of interest (or TSS corresponding to the RNA polymerase RNAP or RNAPII) at an arbitrary location. The overall random search time or mean first passage time (MFPT) *τ*_*U*_ associated with the TF of interest to find its CRM via a combination of 3D and 1D diffusion can be written as follows [9].

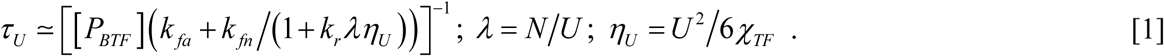

Here *P*_*BTF*_ (M, mols/lit) is the concentration of the TF molecule in the cytoplasm or bulk, *k*_*fa*_ (M^−1^ s^−1^) is the bimolecular rate constant associated with the direct site-specific binding of TF via 3D diffusion route, 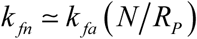 is the overall 3D non-specific binding rate where *R*_*P*_ is the radius of gyration of TF and *k*_*r*_ (s^−1^) is the dissociation rate of nonspecifically bound TF from the DNA lattice. Detailed theoretical studies suggested [9] an expression for *k*_*fa*_ as 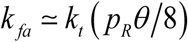. In this expression 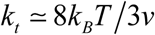 is the maximum possible 3D diffusion limited bimolecular collision rate where *k*_*B*_ is the Boltzmann constant, *T* is the absolute temperature and *v* is the viscosity of the aqueous medium surrounding TFs. The numerical factor 1/8 accounts for the geometry of the random coiled and relaxed conformational state of the DNA polymer. Under *in vitro* laboratory conditions (*T* = 298K and *v* ~ 10^−1^ kg m^−1^ s^−1^ for aqueous solution), one finds that *k*_*t*_ ~ 10^9^ M^−1^s^−1^ [40]. Further *p*_*R*_ is the equilibrium probability of observing a specific or nonspecific binding site to be free from other dynamic roadblock protein molecules [9, 17] which are all present on the same DNA polymer and 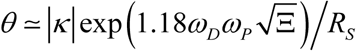 is the factor which account for the overall electrostatic attractive forces and the counteracting shielding effects of the solvent and other ions operating at the DNA-protein interface [9] where *R*_*S*_ ≈ *R*_*P*_ + *R*_*D*_ is the corresponding reaction radius. Here *R*_*D*_ is the radius of the DNA cylinder and 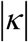 is the Onsager radius which is defined as the distance between the charged reactant molecules at which the overall electrostatic energy will be the same as that of the background thermal energy (i.e. close to 1*k*_*B*_*T*, see **Fig. 1**). Further *ω*_*D*_ and *ω*_*P*_ are the overall charges on the DNA backbone and the DBDs of TFs respectively and Ξ is the ionic strength of the reaction medium.

The term *λ* in **Eq. 1** is the minimum number of association-scan-dissociation cycles required by TFs to scan the entire DNA sequence and *η*_*U*_ is the (averaged over initial positions) overall mean first passage time that is required to scan *U* bps of DNA via 1D diffusion before the dissociation event. Here *U* is a random variable which takes different values in each association-scan-dissociation cycle. The probability density function associated with the 1D diffusion lengths *U* of TFs can be written as follows [9].

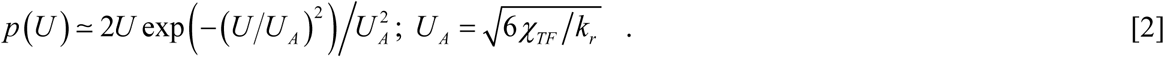

Here *U*_*A*_ is the maximum achievable 1D diffusion length associated with the nonspecifically bound TFs on DNA before the dissociation event that is measured in bps and *χ*_*TF*_ (bps^2^ s^−1^) is the 1D diffusion coefficient associated with the dynamics of TFs. When the TF of interest moves with a jump size of *k* then we find that 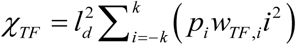 where *i* ∈ *Z*, *w*_*TF,i*_ are the microscopic transition rates associated with the forward and reverse movements of TFs on DNA and *p_±i_* are the corresponding microscopic transition probabilities [8, 41]. Here the step length *i*_*ld*_ is measured in terms of bps where we have defined *l*_*d*_ = 1bps. Since the dynamics at the DNA-protein interface involves segmental motion of DBDs of TF proteins, one can assume the protein folding rate limit [42] for the microscopic transition rates as *w*_*TF*,±1_ ~ 10^6^ s^−1^. Noting that *p_±1_* ~ ½ for an unbiased 1D random walk, one finds that *χ*_*TF*_ ~ 10^6^ bps^2^s^−1^ corresponding to the sliding type dynamics of TFs on DNA for which we have *k* = 1. Approximately this is the experimental value of the 1D diffusion coefficient associated with the sliding dynamics of TFs on DNA [9, 43, 44]. For an arbitrary jump size of *k* with the average microscopic transition probabilities as *p_±i_* = 1/2*k* and average transition rates as 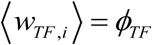, one finds that 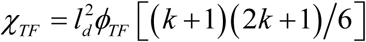. Here one should note that *χ_3D_* will be always ~10-10^2^ times higher than *χ*_*TF*_ and in general we have *χ*_*TF*_ ≤ *χ*_3*D*_ irrespective of the presence of various 1D facilitating processes such as hopping and intersegmental transfers. Various symbols and parameters used throughout this paper are listed in **Table 1**.

**Table 1.** List of symbols and variables used in the theory section

| Symbol | Definition | Remarks |
| --- | --- | --- |
| $x, y$ | Positions of TF and nucleosome on DNA. | m |
| $l_d$ | Length of 1 base-pair (bps) $\sim 3.4 \times 10^{-10}$ m. | m |
| $X, Y$ | $X = x/l_d, Y = y/l_d$ | bps |
| $l$ | Length of DNA lattice under consideration. | m |
| $L$ | $= l/l_d$ | bps |
| $k_{fa}$ | Bimolecular collision rate associated with the formation of site-specific DNA-TF complex via pure 3D diffusion. | $M^{-1}s^{-1}$ |
| $k_{fn}$ | Bimolecular collision rate associated with the formation of nonspecific DNA-TF complex via pure 3D diffusion. | $M^{-1}s^{-1}$ |
| $k_r$ | Dissociation rate associated with the nonspecific DNA-TF complex. | $s^{-1}$ |
| $\tau_s$ | Overall search time that is required by TF to find its cognate site on DNA via a combination of 1D and 3D diffusion routes. | s |
| $\chi_{TF}$ | $= l_d^2 \sum_{i=-1}^1 (i^2 p_i w_{TF,i})$ . 1D diffusion coefficient associated with the dynamics of TFs along DNA. | $\text{bps}^2 s^{-1}$ |
| $\chi_{NU}$ | $= l_d^2 \sum_{i=-1}^1 (i^2 p_i w_{NU,i})$ . 1D diffusion coefficient associated with the sliding dynamics of nucleosomes along DNA. | $\text{bps}^2 s^{-1}$ |
| $\chi_{3D}$ | 3D diffusion coefficient associated with the dynamics of TFs. | $\text{bps}^2 s^{-1}$ |
| $\chi_{TF,S}$ | 1D diffusion coefficient associated with the stationary state of DBDs of TFs. | $\text{bps}^2 s^{-1}$ |
| $\chi_{TF,M}$ | 1D diffusion coefficient associated with the mobile state of DBDs of TFs. | $\text{bps}^2 s^{-1}$ |
| $\chi_{TF,A}$ | $= (\chi_{TF,S} + \chi_{TF,M}) / 2$ | $\text{bps}^2 s^{-1}$ |
| $\chi_{TF,G}$ | $= 2\chi_{TF,S} \chi_{TF,M} / (\chi_{TF,S} + \chi_{TF,M})$ | $\text{bps}^2 s^{-1}$ |
| $D_{TF}$ | $= \chi_{TF} / \phi_{TF} l_d^2$ | dimensionless |
| $D_{NU}$ | $= \chi_{NU} / \phi_{NU} l_d^2$ | dimensionless |
| $\mu$ | Free energy barrier associated with the single bps movement of nucleosome which is achieved mainly by the formation and propagation of thermal induced 1-bps twist defect at the nucleosome-DNA interface. | Dimensionless, measured in $k_B T$. |
| $\gamma$ | Exponent that characterize the steepness of the potential profile which decides the occupancy pattern of nucleosomes along DNA. | $\text{bps}^{-1}$ |
| $\varepsilon$ | $\varepsilon = \phi_{TF} / \phi_{NU}$ | dimensionless |
| $\phi_{TF}$ | $= \langle w_{TF,i} \rangle$ , average rate associated with the unit bps movement of TFs. | $s^{-1}$ |
| $\phi_{NU}$ | $= \langle w_{NU,i} \rangle$ , average rate associated with the unit bps movement of nucleosomes. | $s^{-1}$ |

### 2.1. Effects of nucleosome roadblocks on the TF search time

With this background, we will quantify the effects of slowly diffusing nucleosomes on the overall search dynamics of TFs. First let us consider a linear DNA lattice of *l* m length. Inside this lattice, we assume that a TF molecule and a bulky nucleosome perform 1D random walks with unit base-pair step size (i.e. sliding type dynamics for which *k* = 1) simultaneously (**Fig. 2**). We denote the arbitrary position of TF on the DNA lattice as *x* and the starting position of the nucleosome as *y* where 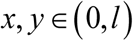. Here the nucleosome span the DNA lattice starting from *y* to *y* + *m*. To simplify the problem, we create a reflecting boundary condition at *x* = 0 and set up the initial positions of TF and nucleosome such that, 0, ≤ *x* < *y* < *l*. The specific binding site corresponding to the TF molecule is located at *x* = *l*. When the length of DNA which is spanned by the bound nucleosome is *m ~* 147 bps, then the specific binding site of TF will be visible to the incoming TF only when the starting point of nucleosome *y* crosses *l*. With this setting the simultaneous 1D diffusion dynamics of TF and nucleosome along the same DNA lattice can be described by the following set of Langevin type stochastic differential equations.

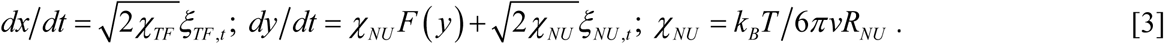

**FIG. 2.**
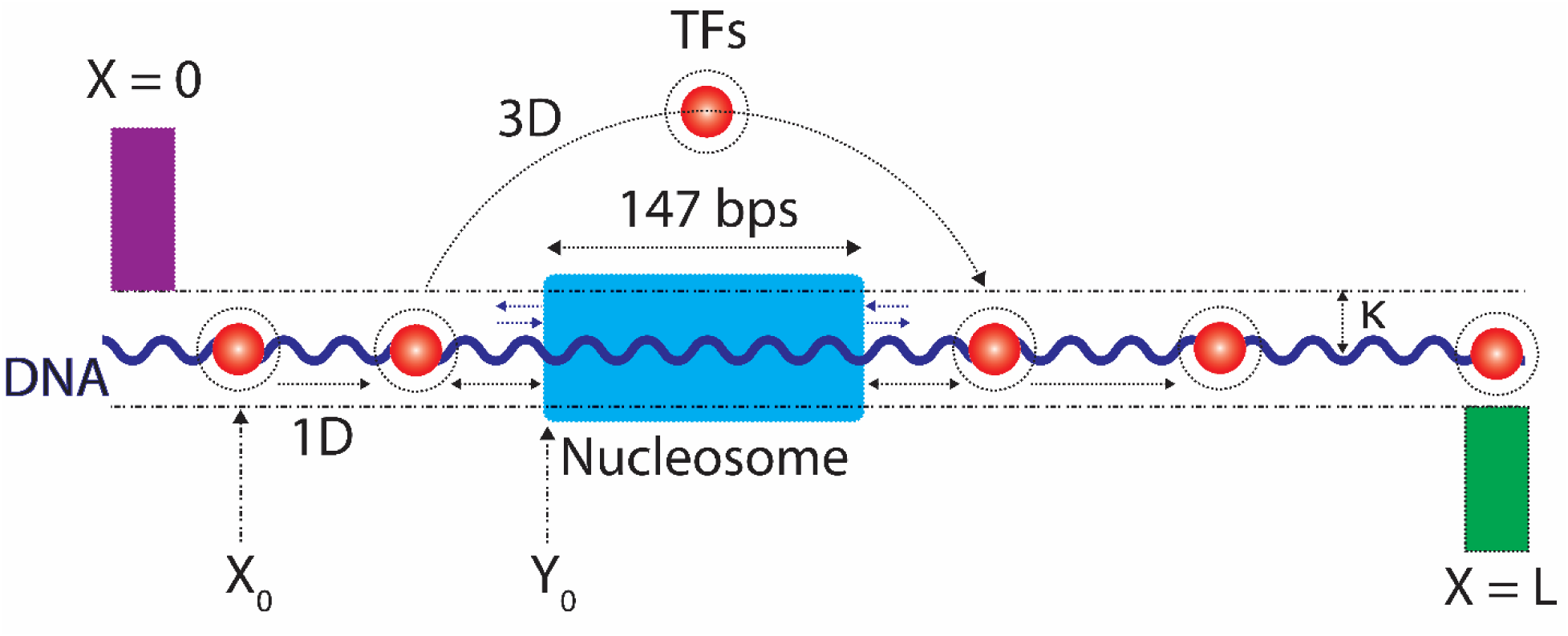
Modelling nucleosome as a massive roadblock across the 1D diffusion of TFs towards their CRMs on the genomic DNA. Here *κ* (bps) is the Onsager radius associated with the electrostatic forces acting at the DNA-protein interface of nonspecific TF-DNA complex. For checking the validity of **Eq. 17** we considered a linear lattice of DNA of *L* bps. We denoted the positions of TF and nucleosome inside this lattice as *X* and *Y* respectively. Actually nucleosome roadblock spans around 147 bps of the DNA lattice which is inaccessible for the inflowing TFs. We introduced a reflecting barrier at *X* = 0 and the CRM of TF is located at *X* = *L*. When TF hits *X* = *L*, then the random walk simulation ends which in turn requires a precondition that *Y* > *L*. However nucleosome can freely diffuse across CRM towards (*L*+1, ∞) and subsequently can also return back into (0, *L*). When the jump size associated with the dynamics of TF is *k* = 1 and *Y* > *X*, then the position of the nucleosome will act as a dynamic reflecting boundary for the movement of TF. When *Y* < *X*, then the dynamic reflections of the nucleosome on TF will enhance the dynamics of TF towards its CRM. When the jump size *k* > 150, then TF can easily jump across the nucleosome and hence the dynamical effects of the slowly diffusing nucleosome on the search dynamics of TF will be at minimum.

Here *χ*_*TF*_ and *χ*_*NU*_ are the 1D diffusion coefficients associated with the dynamics of TF and nucleosome respectively. The force term 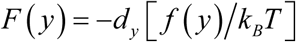 that acts on the 1D diffusion dynamics of nucleosomes is generated by the position dependent dimensionless potential [*f*(*y*)/*k*_*B*_*T*] that is measured in terms of number of *k*_*B*_*T* units. In **Eqs. 3**, *R*_*NU*_ is the radius of gyration of the nucleosome core particle, *v* is the microscopic viscosity of the medium which is surrounding nucleosomes, *k*_*B*_ is the Boltzmann constant and *T* is the absolute temperature in degrees K. The 1D sliding mediated nucleosome repositioning seems to be achieved mainly by the formation of thermal induced bulge defects or twist defect [36] in the interaction network at the DNA-nucleosome interface as shown by Schiessel et.al. in Ref. [39]. Although the repositioning dynamics of nucleosomes shows an anomalous type diffusion under crowded environments [45, 46], we assume a normal type diffusion since we mainly deal here with the interactions between single TF molecule with single nucleosome particle present on the same DNA lattice. We will show in the later sections that even single TF can exhibit a typical sub-diffusive type dynamics especially when it diffuses adjacent to a slowly moving nucleosome particle. Typical nucleosome positioning pattern on the genomic DNA seems to be approximately a periodic one. Here one can consider two different types of potential functions viz. (1) potential function related to the microscopic events occurring at the DNA-nucleosome interface which controls the sliding dynamics of nucleosomes and also decide the value of the 1D diffusion coefficient *χ*_*NU*_ and, (2) sequence dependent potential function related to the observed stationary positioning pattern of nucleosomes along DNA. Beidokhti et.al., in Ref. [37] used a periodic potential of type 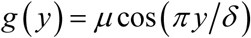 and a three-state model to describe the stochastic sliding dynamics of nucleosomes along DNA. Here the 1D sliding of nucleosome seems to be achieved by the formation of thermal induced 10-bps bulge defect (*δ* = 10) as well as 1-bps twist (*δ* = 1) defect across the interaction network present at the DNA-nucleosome interface and their propagation through the reptation dynamics of DNA [39]. The free energy barrier associated with the formation and propagation of 1-bps twist defect seems to be *μ* ~ 9 *k*_*B*_*T* which is much cheaper than the 10-bps loop defect that involves a free energy barrier of *μ* ~ 23 *k*_*B*_*T* [36, 37, 47]. All these microscopic events and the corresponding free energy barriers will eventually decide the observed value of 1D diffusion coefficient of nucleosome dynamics.

Although the stationary occupancy pattern of nucleosomes seems to be a periodic one, it is still not clear about the existence of periodic type potential along DNA since these are all dynamical entities which occur as a consequence of sequence dependent interaction of nucleosome cores with DNA. Further the distance between the adjacent peaks in the occupancy profile seems to be less than or equal to the length of DNA polymer that is wrapped around the nucleosome particle [13]. Since we are mainly interested in unraveling the interplay between the dynamics of TFs in the presence of nucleosomes, we first develop our model by assuming that both nucleosomes and TFs are unbiased random walkers on the same DNA lattice under constant potential so that the force term in **Eqs. 3** can be set as *F*(*y*) = 0. We will introduce the position dependent occupancy pattern of nucleosomes and the corresponding potential function into our model in the later sections. With this background, in **Eqs. 3** *ξ*_*TF,t*_ and *ξ*_*NU,t*_ are the delta correlated Gaussian white noise terms associated with the stochastic dynamics of TF and nucleosome respectively with the following mean and covariance properties.

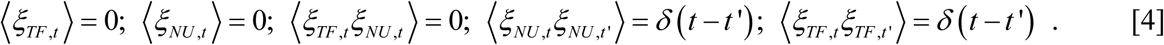

The Fokker-Planck equation (**FPE**) associated with the temporal evolution of the probability of observing TF and nucleosome particle at positions (*x*, *y*) on the same DNA lattice at time *t* with the condition that they were at (*x_0_*, *y_0_*) at *t* = *t_0_* = 0 can be written as follows [41, 48, 49].

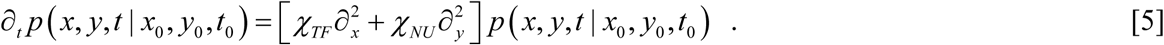

Here initial condition is 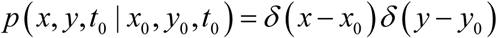 and the boundary conditions for the probability flow can be written as follows.

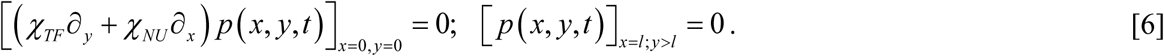

Here both the left side boundary of DNA at *x* = 0 and the position of nucleosome that is present at the right side of TF i.e. *x* = *y* act as reflecting barriers for the dynamics of TFs. In the same way, the TF molecule that is located at the left side of nucleosome i.e. *y* = *x* will be a reflecting boundary for the dynamics of nucleosome. These conditions are required to account for the excluded volume effects i.e. TF and nucleosome cannot occupy the same location on the DNA lattice. Further we assume a natural type or free right side boundary condition for the dynamics of nucleosome. This means that nucleosome can freely cross the location of CRMs and explore further regions (*l*, ∞) of DNA under consideration. To simplify our calculations we carry out the following scaling transformation of the dynamical variables (*x*, *y*, *t*) to make FPE **Eq. 5** in to a dimensionless one.

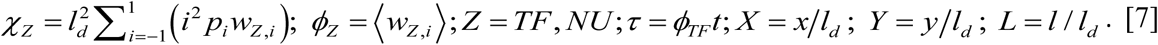

In this equation 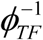 and 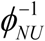 are the average times required for TFs and nucleosome respectively to move unit bps step in the forward and backward direction along DNA. The FPE corresponding to **Eq. 5** can be rewritten in a dimensionless form as follows.

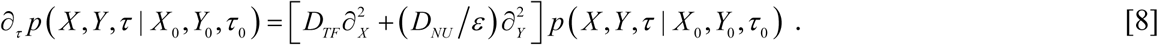

Here the dimensionless 1D diffusion coefficients and other parameters are defined as follows.

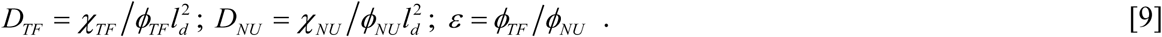

Since the microscopic transitions involved in the 1D diffusion dynamics of TFs require segmental motion of DBDs of TFs, one can assume a protein folding-unfolding rate limit for the unit base-pair displacements along DNA as *ϕ*_*TF*_ ~ 10^6^ s^−1^ from which one obtains *χ*_*TF*_ ~ 10^6^ bps^2^s^−1^. On the other hand the 1D diffusion dynamics of nucleosomes along DNA requires thermal induced formation of bulge or twist defects at the DNA-histone interface and their propagation through corkscrew type dynamics of the DNA polymer [36, 37, 39]. Particularly a free energy barrier of *μ* ~ 9*k*_*B*_*T* is involved in the 1-bps twist driven displacement of nucleosome along DNA [36, 37, 50]. This means that the rate associated with the single bps movement of nucleosomes will be in the order of 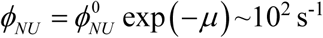 as suggested by the transition state theory (TST) [40] where we have used the protein folding rate limit for 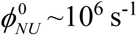 which actually corresponds to a free energy barrier of *μ* = 0. As a result one finds that 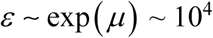 for the barrier *μ* ~ 9*k*_*B*_*T*. Using the value of *ϕ*_*NU*_ one can compute the 1D diffusion coefficient associated with the sliding dynamics of nucleosomes in the absence of other DNA binding ligands and sequence dependent potentials as 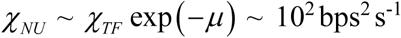 [36]. Clearly we have *ε* = 1 for the roadblock effects of similar types of TFs preforming 1D sliding on the same DNA. With this background, the initial and boundary conditions corresponding to **Eq. 8** are defined as follows [41].

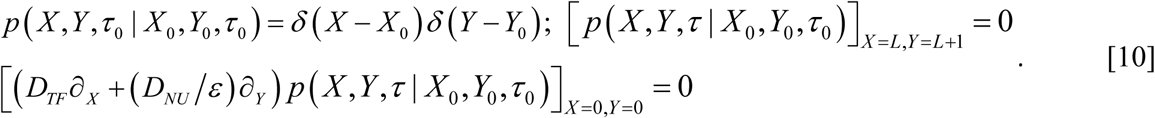

The expression for the mean first passage time (MFPT) associated with the finding of CRMs by the respective TFs in the presence of nucleosome roadblock can be derived as follows. Let us first define the overall probability of observing both TF and nucleosome at (*X*, *Y*) which actually started from (*X_0_*, *Y_0_*) at *τ* = *τ_0_* = 0 still inside the lattice (0, *L*) until an arbitrary time point *τ* as follows.

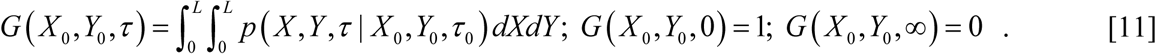

With this definition, **Eq. 8** will transform as follows.

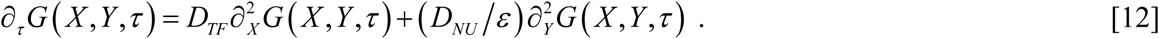

In this equation (*X*, *Y*) are the running variables which represent the initial positions of TF and nucleosome on the DNA lattice respectively and 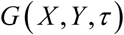 is the probability of observing TF and nucleosome still inside (0, *L*) starting from (*X*, *Y*) until an arbitrary time *τ*. Here one can partition the total amount of time that is required for the binding of TF with its CRM starting from *X* into at least three different time components viz. (1) the time (*τ_1_*) required for the nucleosome to reach CRM that is located at *L* starting from *Y > X*, (2) the time required by the nucleosome (i.e. *τ_2_* – *τ_1_*) to dislocate from the CRM site and (3) the time that is required by TF to reach the already exposed CRM starting from *X* < *Y* via 1D diffusion dynamics along DNA. Clearly *τ*_1_ and *τ*_2_ are random variables and one cannot define the problem of MFPT until the nucleosome completely crosses over the CRM location and subsequently the CRM of TF is exposed for the binding of TF. This is similar to that of the co-transcriptional splicing phenomenon [33] where the small nuclear ribonucleo proteins (snRNPs) need to wait on the pre-mRNA for the emergence of splicing sites (cognate sites of snRNPs) from the transcription assembly to initiate splicing reaction. As a result one obtains the following relationships corresponding to three different time domains of the survival probability function 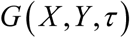.

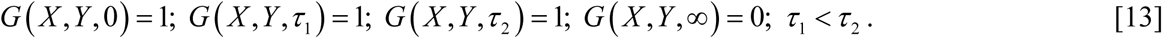

Expressions given in **Eqs. 13** are logical since the probability of finding TF and nucleosome inside the interval (0, *L*) over the timescale (0, *τ_1_*) will be one and so on. From **Eqs. 13** one can derive the following backward type FPE for the overall MFPT associated with the finding of CRM by the respective TF in the presence of nucleosome roadblock starting from the DNA lattice locations (*X*, *Y*) such that *X* < *Y*.

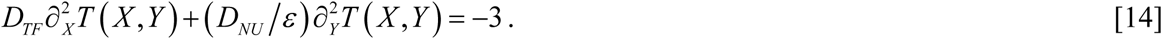

This equation follows from the fact that 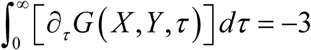 where the overall MFPT which comprises of all the three time domains can be defined as follows.

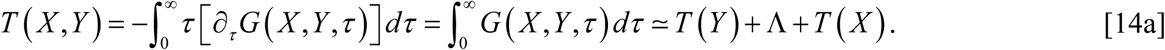

Here various time components in the definition of the overall MFPT are defined as follows.

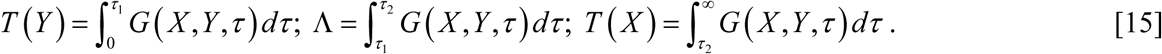

This follows from the fact that 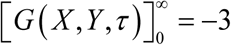 corresponding to three different time domains. The specific boundary conditions for the backward type FPE **Eq. 14** can be written as follows.

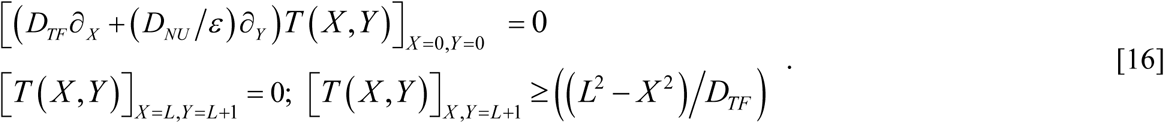

We are introducing an additional boundary condition (third one in **Eqs. 16**) mainly to account for the fact the MFPT should be equal to *T*(*X*) in the absence of nucleosomes. Upon imposing these boundary conditions, one can obtain the following approximate expression for the overall MFPT associated with the finding of CRMs that is located at *L* by TF starting from *X* in the presence of nucleosome roadblock starting from *Y* as follows.

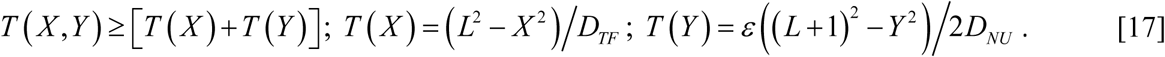

Clearly 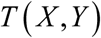 that is defined in **Eqs. 17** is an integral solution of the partial differential equation given in **Eq. 14** which can be straightaway verified by substitution. When 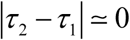 then one can set Λ ≃ 0. This is a reasonable approximation since experimental studies suggested a timescale of ~0.25s for the unwrapping and ~0.05s for the rewrapping dynamics of DNA-bound nucleosome particles [46]. On the other hand the timescale associated with the search dynamics of TFs towards their CRMs spans over several seconds to minutes which proportionately increases with respect to the size of DNA that contains the respective CRM. In this situation **Eq. 17** suggests the following linear type functional relationship between the overall MFPT 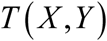 and the delay factor *ε* associated with the dynamics of the nucleosome roadblock.

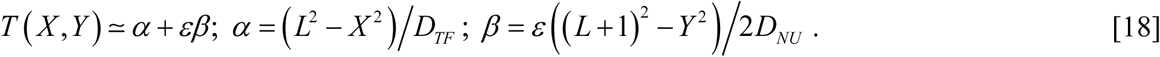

Here *α* is the mean time required by TF to find its CRM via 1D diffusion along DNA which is located at *L* starting from *X* in the absence of nucleosome roadblock. Similarly *β* is the average time required by the nucleosome to cross the CRM of TF located at *L* starting from *Y* in the absence of TF via 1D diffusion. The delay factor *ε* = *ϕ*_*TF*_/*ϕ*_*NU*_ is a critical one which is the measure of the roadblock effects of slowly diffusing nucleosome. While deriving **Eqs. 17-18**, we have assumed the initial condition as 0 < *X* < *Y* < *L* and a jump size of *k* = 1 for the dynamics of TF as well as nucleosome. In the later sections we will show that nucleosomes can enhance the search dynamics of TFs towards their CRMs when their initial conditions are such that 0 < *Y* < *X* < *L*. Especially when 0 < *Y* < *X* < *L* then we will show using simulations that 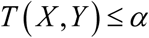 irrespective of the values of *ε*. This enhancement effect over MFPT mainly originates from the dynamic reflections and confinement of the search space of TF on DNA by nucleosome [51]. When *ε* = 0, then the random search dynamics of TFs will not be influenced by the dynamics of nucleosome which is evident from **Eqs. 17-18**. The condition *ε* = 0 will be generally achieved by tweaking the DNA sequence such that nucleosomes dissociate immediately upon contacting with DNA or they diffuse over those sequences under consideration much faster than TFs. When *ε* is very large in magnitude, then the search dynamics of TFs towards their respective CRMs will be exclusively controlled by the nucleosome dynamics.

When *k* > 1 (or *k* > *m* in general where *m* is the length of DNA spanned by nucleosome core) for TF and *k* = 1 for the nucleosome, then the TF molecule can easily jump over the nucleosome roadblocks and therefore the overall search time will not be influenced much by the nucleosome dynamics. At the same time the 1D diffusion coefficient associated with the dynamics of TFs (*D*_*TF*_) will be scaled up as 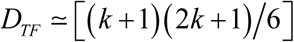. Consequently, irrespective of the values of *ε* (where *ε* > 0) one finds that 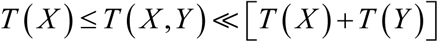. Since the nucleosome complex spans around *m* ~ 147 bps of the eukaryotic DNA, TFs require a jump size of at least *k* ~ 150 bps to completely overcome the roadblock effects of nucleosomes. Here one should note that *k* is positively correlated with the degree of supercoiling or condensation of DNA. Highly condensed conformational state of DNA favors higher jump sizes of TFs. On the other hand linear or stretched conformational state of DNA always favors sliding type dynamics TFs. To understand the exact roles of nucleosomes on the dynamics of TFs one can consider the initial position averaged MFPT in terms of the original variables as follows.

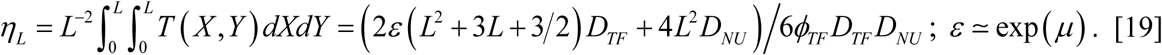

In this equation *η*_*L*_ (s) is the average time required by a TF molecule to scan *L* number of sites on DNA in the presence of a nucleosome where we have the limit 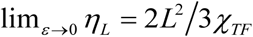. Because of the asymmetric type dynamics of TFs in the presence of nucleosomes and there is a reflecting boundary at *X* = 0, *η*_*L*_ is four times more than *η*_*U*_ of **Eqs. 1**. When *ε* is sufficiently large then one obtains 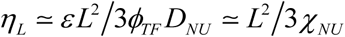. In general one finds that 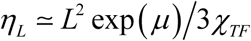 where we have used the approximation 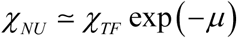. **Eqs. 19** suggest that the effective 1D diffusion coefficient associated with the search dynamics of TFs towards their cognate sites will be decreased in the presence of nucleosomes and subsequently 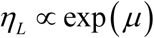. These results suggest that the relative amount of time spent by TFs on 1D and 3D diffusion routes can be fine-tuned by the slow dynamics of nucleosomes.

### 2.2. Effects of nucleosome occupancy pattern on MFPT

It is remarkable to note that the nucleosome-depleted regions of the genomic DNA correlate well with the transcription start sites (TSSs) as well as transcription factor binding sites (TFBSs) or CRMs on genome wide level [30]. Here the nucleosome depleted regions observed on the genomic DNA can occur in two possible ways viz. (1) nucleosomes cannot bind with those DNA sequences at all and (2) the delay factor *ε* is gradually decreasing in a sequence dependent manner while nucleosomes approach the locations of TSS and TFBS. As a result of these facts, nucleosomes undergo large extent of breathing type dynamics as they approach and cross over the regions of TSS and TFBS [28]. In contrast, the extent of conformational fluctuations in the DBDs of TFs attenuates as they move close to their CRMs [19]. These results suggest that the parameter *ε* will be strongly dependent on the distance of the current position of nucleosome from TSSs and TFBSs. In other words, one needs to apply certain distance dependent weightage over *ε* rather than a uniform one. Here the weightage term could be the normalized occupancy pattern of nucleosomes for a given stretch of DNA sequence where the occupancy value is directly proportional to the probability of observing nucleosomes at a given position on DNA. Close observation over *in vivo* MNase-seq data on the nucleosome occupancy pattern around TSSs and CRMs suggests the following functional form [34, 52].

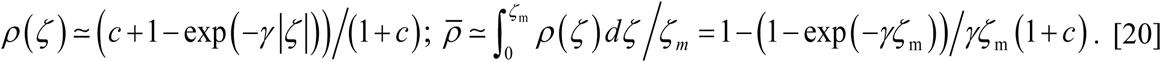

In this equation *ζ = L − Y* (bps) is the distance of the location of nucleosome from TSSs and CRMs (that is located at *L* in the present context so that *ζ* = 0 corresponds to the location of TSS and CRMs and the maximum distance will be *ζ*_*m*_ = *L*) and *γ* (bps^−1^) is the critical exponent which is inversely proportional to the steepness of the nucleosome depletion valley that is surrounding the locations of TSS and CRMs. Here 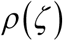 is the normalized distance dependent nucleosome occupancy which in turn modulates the values of *ε*. The value of 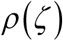 at a given location will be directly proportional to the probability of observing a nucleosome particle at that location. The normalized average nucleosome occupancy around the location of TSSs and CRMs is defined as 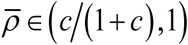 where the constant *c* is the genome wide average of the nucleosome occupancies at the locations of TSSs and CRMs i.e. average value of the minimum nucleosome occupancy at TSSs and TFBSs. The boundary values of 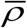 will be dictated by the following limiting conditions.

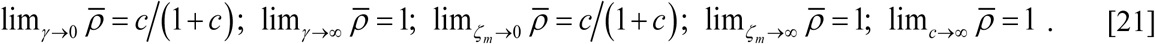

Upon considering the position dependent weightage for the nucleosome occupancy pattern around TSS and CRMs, one can obtain the overall expression for the MFPT associated with the finding of CRMs by TFs in the presence of nucleosome roadblocks as 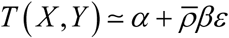. This clearly suggests that the roadblock effects of nucleosomes can be nullified by properly tuning their sequence dependent occupancy pattern. This is evident from the fact that when 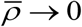 (which can be obtained in the limit as *c* → 0) then the value of the MFPT will be close to the one that is obtained in the absence of nucleosomes. Here one should note that presence of significant amount of site-specific DNA-TF interactions around these TSSs or TFBS could also be one of the reasons for such nucleosome-depletion apart from the weak DNA-nucleosome interactions. Actually nucleosomes need to compete for the freely available DNA segments around TSSs and TFBSs with other TFs and protein factors which are involved in the transcription initiation. Detailed analysis on the nucleosome occupancy pattern around the CRMs of various TFs of human genome shows a minimum around the regions of CRMs with well positioned nucleosomes around such minimum occupancy locations [29]. It is interesting to note that the normalized occupancy pattern of TFs shows a maximum at CRMs and one can use the following type function as in **Eq. 20** to model such occupancy pattern of TFs.

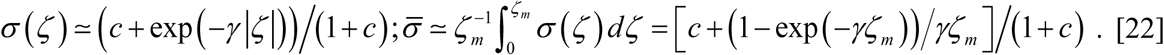

Here 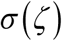 is the distance dependent occupancy of TF on the DNA lattice where *ζ = L − X* is the distance of TF from its CRM location, *c* is the background TF occupancy corresponding to the genomic DNA and *γ* is the exponent which characterizes the shape of the occupancy profile. The maximum possible distance in the present context will be *ζ*_*m*_ = *L*. The overall average of the occupancy of TF over CRMs will be given by 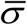 of **Eqs. 22** with the following limiting conditions.

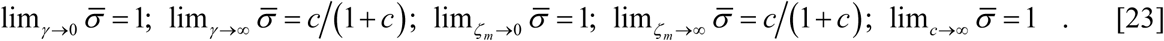

Here one should note that 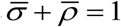. The occupancy profile of nucleosomes around the distal TFBS under *in vitro* conditions as well as over the repressor binding regions of the genomic DNA under *in vivo* conditions seems to be similar to that of 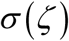 as in **Eqs. 22**. Under such conditions the overall MFPT associated with TFs to find their respective CRMs in the presence of nucleosome roadblocks will be scaled up as 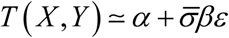 where *α* and *β* are defined as in **Eqs. 18**. The main drawback of the occupancy weighting functions 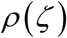 and 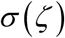 is that they do not consider the underlying periodic behavior of the occupancy profiles of nucleosomes [13] though the amplitude of such oscillations in the vicinity of TSS or TFBSs seems to be negligible [52].

### 2.3. Dynamics of nucleosomes in the presence of position dependent potential

One can reverse calculate the probable potential functions which can give rise to the observed stationary probability density function similar to that of the occupancy profile of nucleosomes along the genomic DNA under *in vivo* conditions. In this context, the FPE corresponding to an arbitrary force function *F*(*y*) can be written as follows.

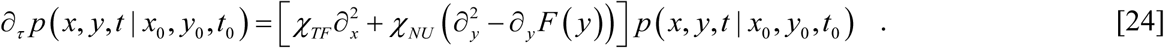

Here the force term is defined as 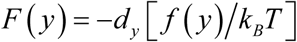. The potential function 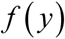 in **Eq. 24** with a period of ~100 bps [13, 52] arises as a consequence of sequence dependent interactions of nucleosome core particles with DNA. Clearly 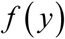 decides the stationary state occupancy pattern of nucleosomes over the genomic DNA as given in **Eqs. 20** and **22** and it is different from the potential function associated with the unit-bps sliding dynamics of nucleosome with period of ~10 bps [37]. The initial and boundary conditions corresponding to **Eq. 24** are similar to that of **Eqs. 5** and **6**. The corresponding FPE in the dimensionless form can be derived as follows.

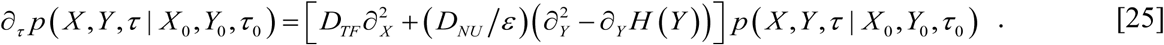

In this equation the dimensionless force term is defined as 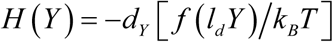. The initial and boundary conditions corresponding to **Eq. 25** are similar to that of **Eqs. 10**. The backward type FPE which describes the MFPT associated with the finding of CRM by TF in the presence of nucleosome roadblock under an arbitrary potential can be written as follows.

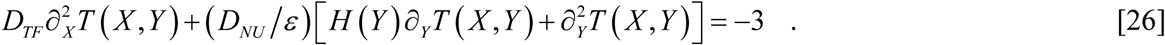

The integral solution of this backward FPE can be written as a sum of three time components as in the case of **Eq. 14**. Various time components associated with the integral solution of **Eq. 26** 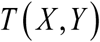 can be written explicitly as follows.

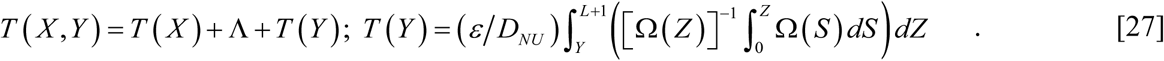

Here we have defined the function 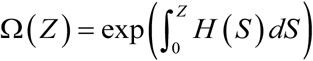 [41]. **Eq. 27** is the particular solution of **Eq. 26** which can be checked by substitution. Beidokhti et.al., in Ref. [37] suggested a periodic type potential as 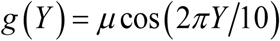 with a period of 10 bps and *µ* ~ 9*k*_*B*_*T* (corresponding to 5S sea urchin positioning element) [36] to describe the microscopic details of 1D sliding dynamics of nucleosomes along DNA. We have already included the effect of this microscopic barrier in to the definitions of the delay factor *ε* as 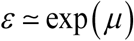 and the 1D diffusion coefficient associated with the dynamics of nucleosomes along DNA as 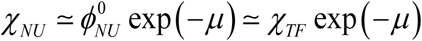. We propose here a position dependent interaction potential of type 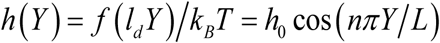 to describe the stationary state occupancy pattern of nucleosomes along DNA of length *L* bps. The force generated by this potential will be 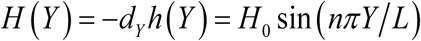. Remarkably this potential function can generate a periodic type stationary occupancy profile of nucleosomes across the DNA sequence under consideration. Upon considering the sequence information, the value of *n* in the expression for 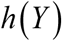 needs to be properly adjusted to reverse calculate the probable potential function which can give rise to the observed occupancy profile of nucleosomes along DNA. For example, to generate an occupancy profile with 6 and 10 peaks per 1000 bps, one needs to set *n* to 12 and 20 respectively. The stationary distribution function 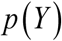 associated with the nucleosome occupancy profile on DNA can be written as follows [41].

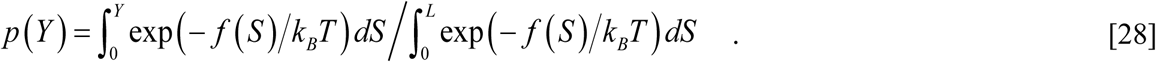

In the derivation of **Eq. 28** we have assumed a reflecting boundary conditions at *Y* = *L* and *Y* = 0 for the dynamics of nucleosomes inside (0, *L*). In the expression of the potential function 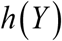, the term *h*_0_ is the dimensionless amplitude which is measured in terms of number of *k*_*B*_*T* units. Since the periodic function described here can go from –*h_0_* to +*h_0_* over the energy axis, the maximum peak to trough distance of 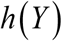 will be 2*h_0_*. This is approximately the height of the free energy barrier associated with the transition of a nucleosome particle from one minimum to another adjacent one over the occupancy profile along DNA. This free energy barrier needs to be fine-tuned for an efficient nucleosome positioning as well as sliding across each local minimum. Particularly high level of free energy barrier leads to strong positioning of nucleosomes but their capability to slide through a minimum will be very much limited.

On the other hand, low barrier heights leads to fast sliding across local minima but poor positioning and repositioning. Our earlier studies suggested that the efficiency of thermodynamic coupling [20, 53] between the extent of conformational fluctuations in the DBDs of TFs and their random 1D search dynamics along DNA will be at maximum only when the free energy barrier associated with the conformational fluctuations in the DBDs of TFs is <3*k*_*B*_*T*. This is similar to that of the dynamics of downhill folding proteins at their mid-point denaturation temperatures [54]. These results further suggest us a condition to achieve the maximum efficiency of nucleosome positioning-repositioning along DNA as 2*h_0_ <* 3. This means that *h_0_ <* 3/2 *k*_*B*_*T*. For the purpose of numerical calculations we set *h_0_* = 1. When the potential function is a constant one along DNA then the force generated by such constant potential will be 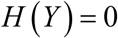. Under such conditions we recover our earlier expression for the MFPT that is described in **Eqs. 17** and **18**. To model the nucleosome depleted basin around TSSs and TFBSs one can use a potential function such as 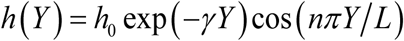 with asymptotically decreasing amplitude. Upon properly setting the value of *n*, *γ*, and *h_0_* one can generate the occupancy profiles with fine details similar to 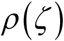 and 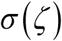 where *ζ* = *L − Y*.

### 2.4. Effect of nucleosomes on the overall search time of TFs

Upon considering the factors viz. (1) the free energy barrier (*μ*) associated with the microscopic transitions involved in the 1D sliding dynamics nucleosomes as given in **Eqs. 19** and the (2) sequence dependent potential which give rise to the observed occupancy profile of nucleosomes along DNA as given in **Eqs. 20** and **22**, the expression corresponding to the overall search time (*τ*_*S*_) given in **Eq. 1** can be rewritten as follows.

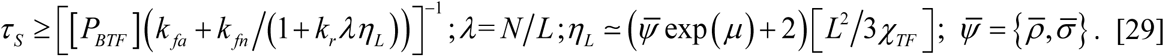

**Eq. 29** suggest that lim_*µ*→∞_ *η*_*L*_ = ∞ and subsequently 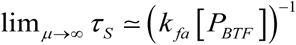 which is exactly the time that is required by TFs to locate their cognate sites on DNA via pure 3D diffusion routes. This also means that 1D diffusion is not an efficient route for random searching of TFs over large eukaryotic genomes. We are using an inequality relationship here mainly due the fact that **Eq. 29** ignores the occurrence of sub-diffusive dynamics of TFs when they move close to the nucleosome roadblocks. Interestingly **Eqs. 29** also suggest that the roadblock effects of nucleosomes can be minimized upon setting 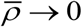 i.e. presence of nucleosome depleted regions around TFBS plays critical roles in minimizing the roadblock effects in those regions. On the other hand, strong positioning pattern and slow sliding dynamics of nucleosome over other regions of genomic DNA ensures that TFs are not permanently adsorbed over noncoding DNA via nonspecific binding.

## 3. Simulation results

To check the validity of **Eqs. 17-23** we performed detailed stochastic random walk simulations at various jump size *k* values. The results are summarized in **Figs. 3**-**6**. We defined a linear DNA lattice of *L* = 100 bps in length. We denoted the arbitrary location of TF inside this lattice as *X* and the location of nucleosome as *Y*. Both TF and nucleosome were moved with unit base-pair step size (sliding) i.e. we first considered the jump size *k* = 1. This means that *D*_*TF*_ =1, and *D*_*NU*_ = 1. Here *X* = 0 is the static reflecting boundary for the dynamics of both TF and nucleosome and *X* = *Y* will be a dynamic reflecting boundary for TF as well as nucleosome. We set the initial position of TF arbitrarily at *X_0_* = 50 and iterated the initial position of the nucleosome from *Y_0_* = 1 to *Y_0_* = 101. Further TF and nucleosome cannot occupy the same location of the DNA lattice due to the excluded volume effects. We set the location of CRM of the TF molecule at *X* = 100. The forward and backward movements of TF and nucleosome were decided by calling a random number which is equally distributed within (0, 1) i.e. 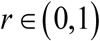 with a probability density function as 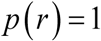. When *r* > 0.5, then the random walker moves forward and when *r* < 0.5 then the random walker moves backward. The possibility of each movement will be decided by the condition that there is no TF or nucleosome at the new location. The delay factor *ε* was introduced into our simulation as follows. For each simulation step, TF can move forward or backward with equal probabilities. However nucleosome can make such forward or backward move only after *ε* number of TF movements. When TF hits the absorbing point then the simulation will be stopped there. However this requires that *Y* > 100 i.e. the nucleosome has already crossed the location of CRM and currently inside 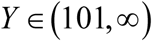. When nucleosome reaches the location *Y* = 101 then it stalls there permanently as per the third boundary condition of **Eqs. 16**.

**FIG. 3.**
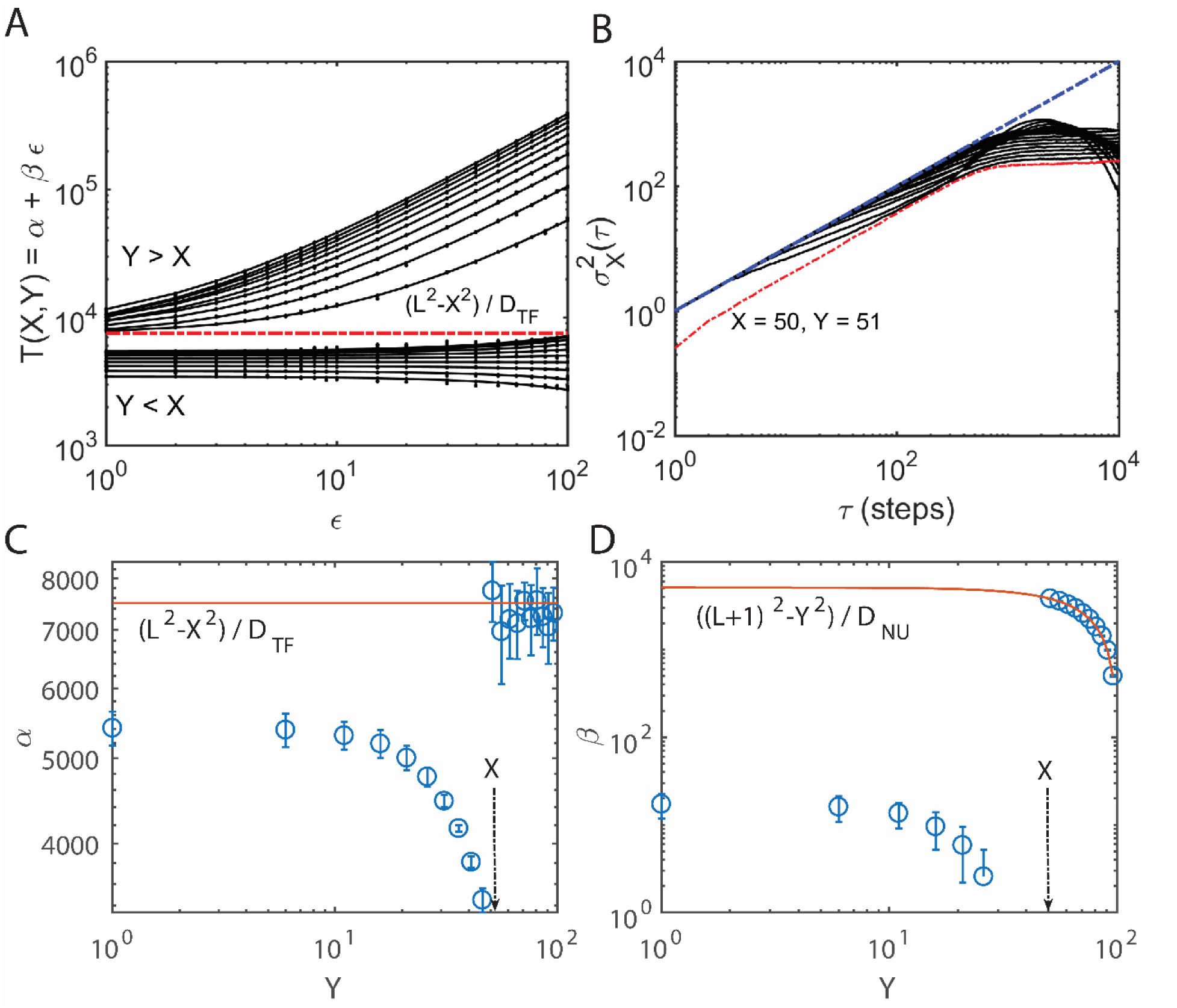
Random walk simulations with jump size *k* = 1 for which *D*_*TF*_ = 1 and *D*_*NU*_ = 1. All these simulations were done on a lattice of size *L* = 100 with reflecting boundary at *X* = 0 and absorbing boundary at *X* = 100. Initial position of TF was fixed at *X_0_* = 50 and the initial position of nucleosome *Y* was iterated from 1 to 100. Each iteration was repeated at different values of *ε* where *ε* was iterated from 1 to 100. The mean first passage time (in terms of number of simulation steps) was calculated over 10^5^ trajectories. Hence obtained MFPT data was used for linear least square fitting to obtain *α* and *β* at a confidence level of 0.95. Those obtained values of *α* and *β* were plotted with respect to *Y* values. **A**. Variation of MFPT with respect to changes in *Y* and *ε*. When *Y* > *X_0_*, then the MFPT data fits linearly with *ε*. When *Y* < *X_0_* then the nucleosome dynamics favors the search dynamics of TFs towards their CRMs due to the dynamic reflections. This is evident from the decreasing values of the intercept *α* (**C**). The degree such enhancement effects increases as the initial distance between TF and nucleosome i.e. |*X* - *Y*| decreases. At the same time this enhancement effect is not dependent of *ε* much which is evident from the values of the slope *β* (**D**). **B**. Variation of the displacement variance with respect to simulation steps. Clearly when *X_0_* = 50 and *Y* = 51, then the system shows a typical sub-diffusion pattern. **C**. Here the settings are *L* = 100, initial *X_0_* = 50 and *DTF* = 1. **D**. Here the settings are *L* = 100, *D*_*NU*_ = 1 and initial *X_0_* = 50. Further *ε* was iterated from 1 to 100.

**FIG. 4.**
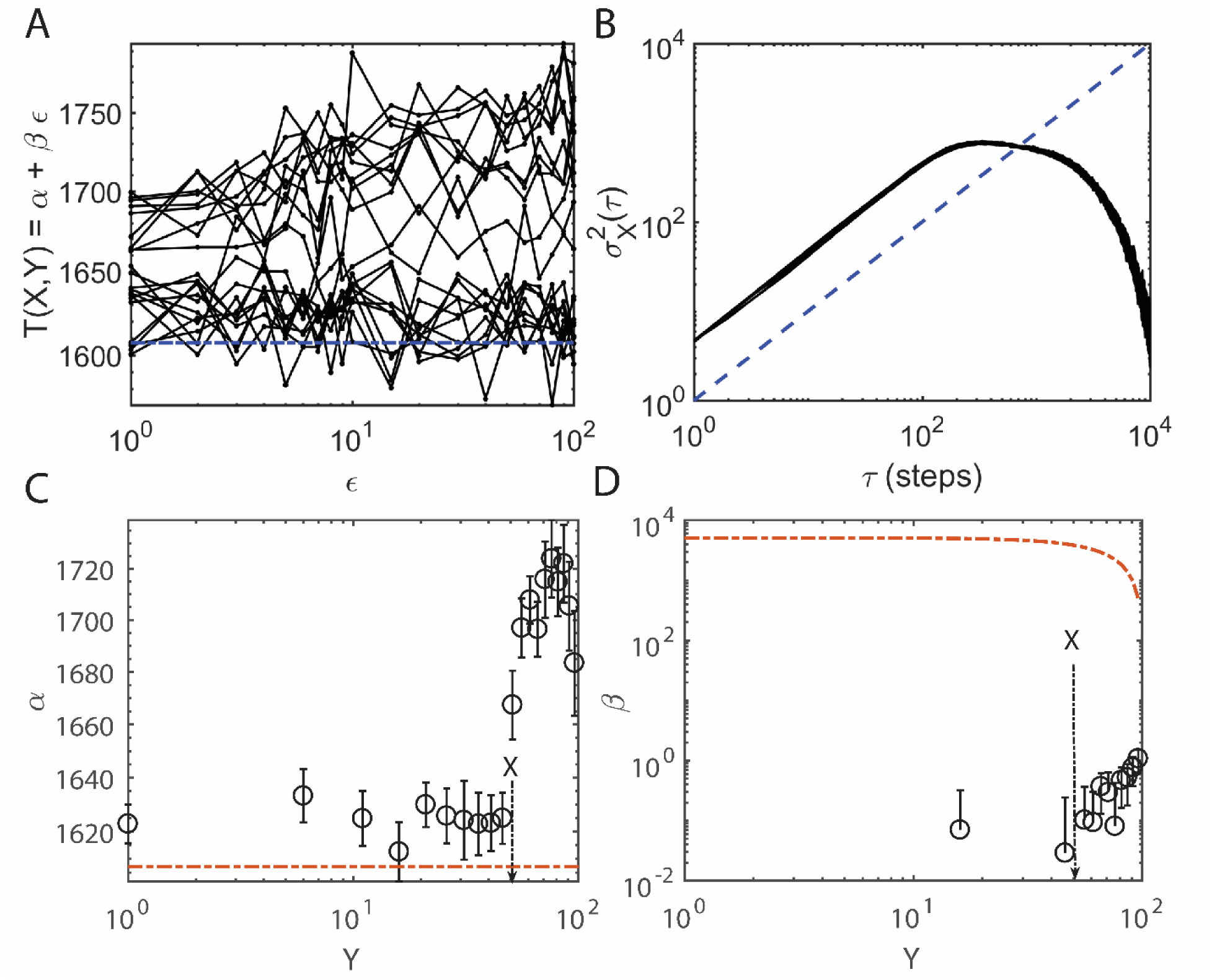
Random walk simulations with jump size *k* = 3. All the random walk simulations were done on a lattice of size *L* = 100 with reflecting boundary at *X* = 0, absorbing boundary at *X* = 100 and initial *X* = 50. The hop size *k* was set to 3 so that the overall multiplication factor for the diffusion coefficient *D*_*TF*_ will be [(*k*+1) (2*k*+1) / 6] = 28/6. Here *D*_*NU*_ = 1 since the jump size for nucleosome is *k* = 1. Since the nucleosome occupies only one lattice location in the present setting, TF can jump across the nucleosome when *k* > 1. Nucleosome initial position *Y_0_* was iterated from 1 to 100. The mean first passage time (in terms of number of simulation steps) was calculated over 10^5^ trajectories at different *ε* from 1 to 100. This MFPT at various *ε* values was used for linear least square fitting to obtain α and β values at a confidence level of 0.95 (**C** and **D**). **A**. Here the settings are *L* = 100, *X* = 50 and *D*_*TF*_ = 28 / 6 and 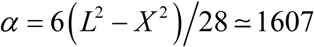 steps (dotted blue line in **A** and dotted red line **C**). **B**. Changes in the variance of TF position *X* with respect to the simulation steps and *Y*. When *k* > 1, then irrespective of whether *Y < X* or *Y > X*, the system show normal diffusion pattern. **C**. Changes in *α* with respect to changes in the position of nucleosome *Y*. **D**. Changes in the value of *β* with respect to changes in the values of *Y*. Clearly when *k* = 3, then the slope *β* seems to be almost independent on *Y*.

**FIG. 5.**
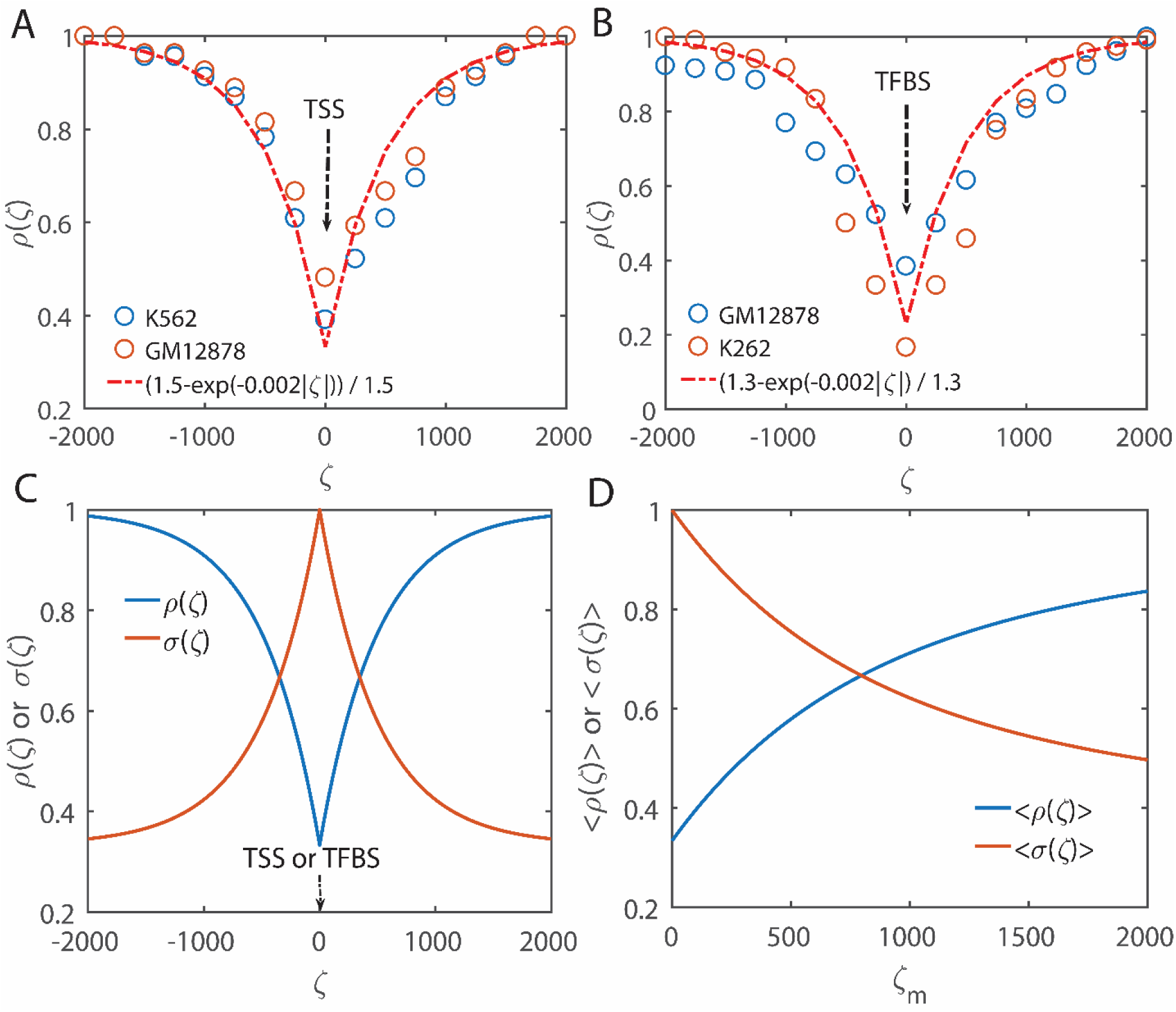
Nucleosome positioning or occupancy pattern around transcription start sites (TSS) and transcription factor binding sites (TFBS, especially proximal) in K562 and GM12878 cell lines under *in vivo* conditions. These plots are based on the MNase-seq data. The normalized occupancy data from Refs. [34, 52] (hollow circles, normalization was done by dividing the digitized data from Ref. [52] with the maximum value of the nucleosome occupancy in the range from -2000 to +2000 bps) seems to follow a functional relationship as 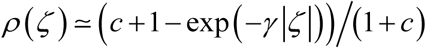 where *ζ* is the distance of nucleosome from TSS (**A**) or TFBS (**B**) on DNA measured in bps, *γ* ~ 0.002 bps^−1^ and *c* ~ 0.5 for the nucleosome occupancy around TSS and *c* ~ 0.3 for the nucleosome occupancy around TFBS. **C**. Theoretical nucleosome occupancies around TFBS and TSSs (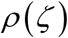) locations and repressor binding sites 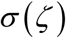 as in **Eqs. 20** and **22**. Settings are *γ* ~ 0.002 bps^−1^ and *c* ~ 0.5. Here the occupancy values are normalized such that the genome wide maximum occupancy is 1. **D**. Variation of positional averages of 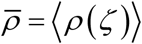 and 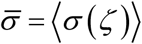 over *ζ* from 0 to maximum *ζ*_*m*_ with respect to *ζ*_*m*_. Settings are *γ* ~ 0.002 bps^−1^ and ĉ 0.5.

**FIG. 6.**
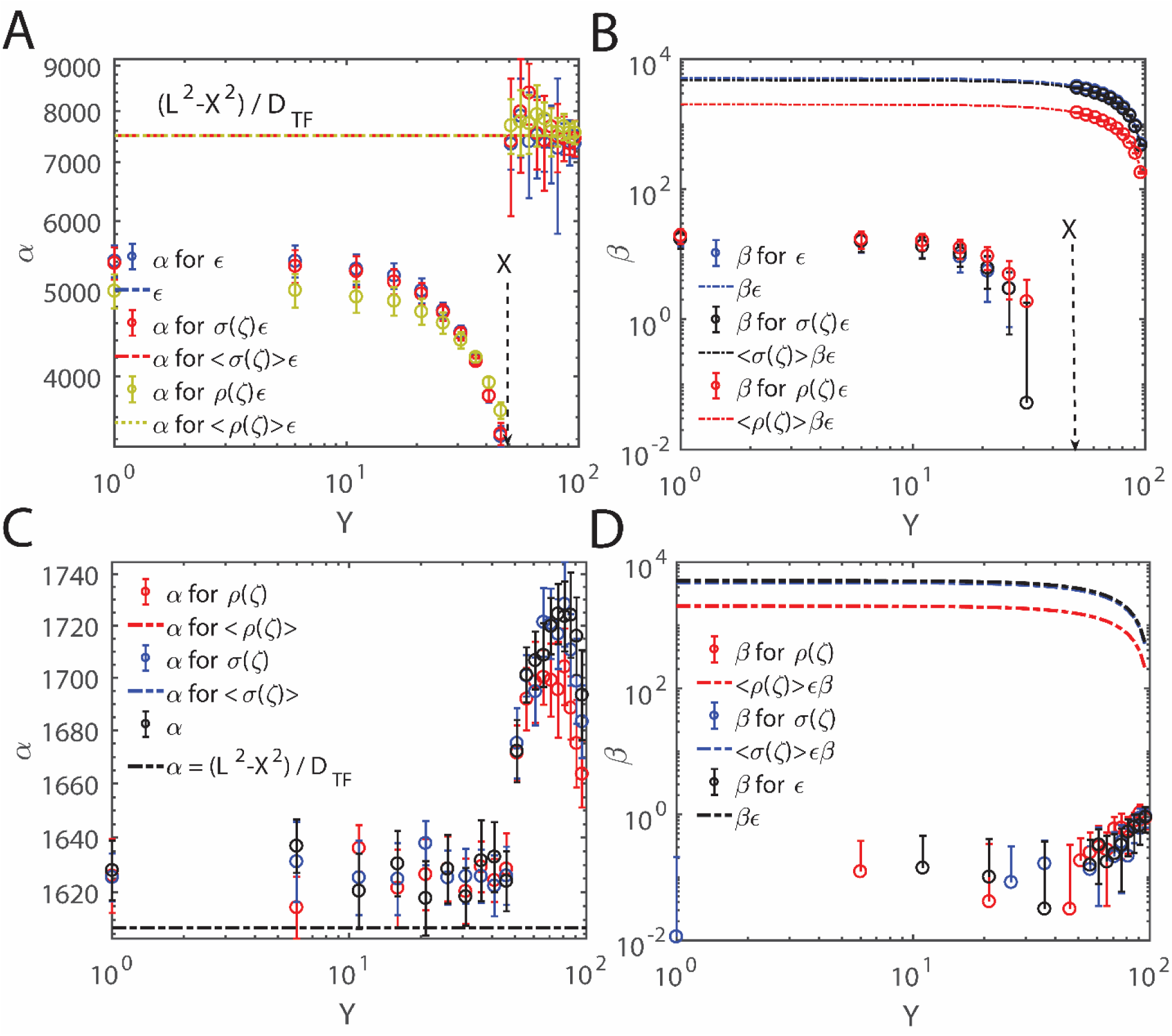
Random walk simulations in the presence of position dependent nucleosome occupancy patterns 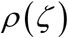 and 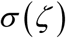 along DNA with settings as *γ* ~ 0.002 bps^−1^ and ĉ 0.5. The overall MFPT seems to be linearly related with respect to *ε* which is modified in the presence of position dependent nucleosome occupancy as 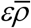 or 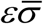 as in **Eqs. 20** and **22**. Here we have defined 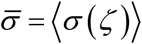 and 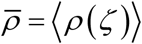 where *ζ* (bps) is the distance between the current position of nucleosome from the location of CRMs and TSSs. Here settings are *L* = 100 with reflecting boundary at *X* = 0 and absorbing boundary at *X* = 100. Initial position of TF was fixed at *X_0_* = 50 and initial position of the nucleosome *Y_0_* was iterated from 1 to 100. Each iteration was repeated at different values of *ε* where *ε* was iterated from 1 to 100. At each simulation step, *ε* was modified as 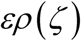 or 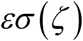 where *ζ = L − Y* as defined in **Eqs. 20** and **22**. The mean first passage time (in terms of number of simulation steps) was calculated over 10^5^ trajectories. Hence obtained MFPT data was used for linear least square fitting to obtain *α* and *β* at a confidence level of 0.95. Those obtained values of *α* (**A** and **C**, for *k* = 1) and *β* (**B** and **D** for *k* = 3) were plotted with respect to the initial *Y* values. Theoretical estimate for *α* is *α* = *L*^2^ − *X*^2^ = 7500 for *k* = 1 (dotted lines in **A**), and it will be 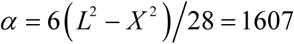 for *k* = 3 (dotted lines in **C**). When *Y* > *X*, then the MFPT data fits linearly with *ε*. However when *Y* < *X*, then the nucleosome dynamics favors the search dynamics of TFs towards their CRMs due to the dynamic reflections and confinement. This is evident from the decreasing values of the intercept *α* (**C**). The degree such enhancement effects increases as the initial distance between TF and nucleosome i.e. |*X* - *Y*| decreases. At the same time this enhancement effect is not dependent of *ε* much (especially when *k* = 3) which evident from the values of the slope *β* (**B** and **D**).

The delay factor *ε* was iterated from 1 to 100 for each value of *Y_0_*. This dataset was used for linear least square fitting procedures to obtain *α* and *β* at a confidence level of 0.95 (error bars) [55, 56]. These are depicted in **Figs. 3C** and **D**. Here *ε* = 0 represents an immediate dissociation of nucleosome roadblock upon contacting with DNA and *ε* = 1 represents TFs like diffusion dynamics of nucleosomes. For example when *ε* = 0 and jump size *k* = 1, then one finds from **Eq. 18** that 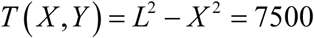 (**Fig. 3C**). Likewise when *ε* = 0 and *k* = 3, then one can compute the 1D scanning time as 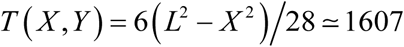 (**Fig. 4C**). In these calculations we have substituted *L* = 100, *X* = 50 and *k* = 1. **Fig. 4C** suggests that 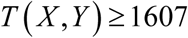 steps for *k* = 3 even at *ε* = 0 which is mainly due to the dynamic reflections of TF upon encountering nucleosome. These stochastic random walk simulation results are in fact consistent with our theoretical predictions given by **Eqs. 17** and **18** especially when *Y* > *X*. The third boundary condition in **Eq. 16** works very well (or true) only when the 1D diffusion dynamics of nucleosome is much slower than that of the TF molecule i.e. at higher *ε* values. This condition will ensure that the nucleosome which has already crossed the location of CRM and currently present inside 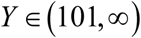, will take much extended time to return back into the interval 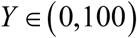. This guarantees a roadblock free environment for the dynamics of TF along DNA after the elapse of *εβ* amount of time. When the speed of nucleosome dynamics is comparable with that of the dynamics of TF (for example *ε* ~ 1) then the nucleosome which has already crossed the CRM (i.e. *Y* > *L*) can return back into the lattice interval 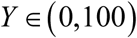 before the site-specific binding of TF with its exposed CRM. Such events will impede the dynamics of TF towards its CRM further. This means that one needs to replace the first equation of **Eqs. 18** with an inequality as 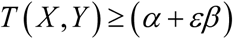.

Here one should note that **Eqs. 17-23** will be valid only when 0 *X < Y < L*. When TF is located in between nucleosome and CRM i.e. 0 *X < Y < L*, then the search dynamics of TF towards its CRM seems to be enhanced irrespective of the values of *ε*. This is evident from the observation that 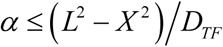 and *β < L* in **Fig. 3C**. However the overall enhancing effects observed when *X* > *Y* are much lesser than the retarding effects of nucleosome when *X* < *Y* as we have demonstrated in our earlier works [51]. It is remarkable to note that when the nucleosome present initially at *Y* = *X* + 1, then the dynamics of TF exhibits a typical sub-diffusive [57] type pattern which is evident from the plot of variance of the position of TF *X* with respect to the number of simulation steps (**Fig. 3B**). Such anomalous type diffusion disappears when the initial distance between TF and nucleosome on the DNA lattice increases or the jump size associated with the dynamics of TF is such that *k* > 1.

### 3.1. Effects of nucleosome occupancy pattern on the TF search time

Here one should note that all these random walk simulations assumed an unbiased weightage on *ε* over the distances between nucleosome and TF from the CRM location. Detailed analysis of the positioning of TFs and nucleosomes along the genomic DNA suggested a biased weightage with respect to the distances of TFs and nucleosomes from the corresponding CRMs. For example, the *in vivo* nucleosome occupancy profile over transcription start sites (TSSs) shows a typical minimum occupancy at TSSs which increases as the distance from the location of TSSs increases [30, 34, 46, 52] towards both upstream and downstream directions. In the following section we will try to understand and unravel the effects of such position dependent dynamics of TFs and nucleosomes on the overall search time.

To understand the effects of position dependent nucleosome occupancy pattern on the TF search time we first computed the normalized nucleosome occupancy values from the MNase-seq data corresponding to the regions surrounding TSS and proximal TFBS. These datasets were actually obtained from K562 and GM12878 cell lines [34]. We first digitized the data from Ref. [52] and normalized it by dividing with the maximum occupancy value occurred inside the window 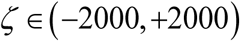. Close observation of such normalized datasets from Refs. [34, 52] suggested a functional form for the position dependent nucleosome occupancy pattern similar to 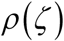 under *in vivo* conditions as in **Eqs. 20**. Numerical simulations of 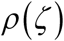 over a wide range of parametric space were carried out to match with the observed occupancy profile. Analysis results suggested the parametric values as *γ* ~ 0.002 bps^−1^, *c* ~ 0.5 for the nucleosome occupancy around TSSs and, *c* ~ 0.3 for the nucleosome occupancy around proximal TFBS (**Fig. 5)**. Here *ζ* is the distance between the position of nucleosome and location of TSS and TFBS on DNA. To include the effects of nucleosome occupancy pattern in the random walk simulations, we modified the value of the delay parameter *ε* of nucleosome at each simulation step in a position dependent manner such that 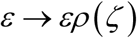 as defined in **Eqs. 20** where *ζ* = *L − Y* is the distance of nucleosome from CRMs. Here *ζ* < 0 denotes the downstream region, *ζ* > 0 denotes the upstream region and *ζ* = 0 denotes the exact location of TSS and TFBS.

The simulation results corresponding to the weighting function 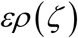 for *k* = 1 are shown in **Figs. 6A-B**. With these settings, our simulation results suggest that the *ε* term in the expression for the MFPT that is given in **Eqs. 18** transform as 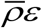 where 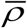 is the overall average of the nucleosome occupancy values around TSS and TFBS as defined in **Eqs. 20**. This in turn reduces the overall MFPT required by TFs to locate their CRMs in the presence of nucleosome roadblocks as 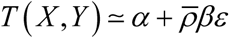 (**Figs. 6A-B**) where we have 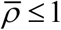. When the distribution of nucleosomes around the repressor binding sites follows a pattern similar to 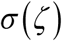 of **Eq. 22**, then the simulation results which are shown in **Figs. 6A-B** agree well with the prediction of the MFPT expression 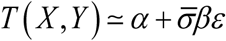. However these approximate relationships seem to break down when *k* > 1. For example, the simulation results corresponding to *k* = 3 are shown in **Figs. 6C-D**. These results suggest that when *k* > 1 then irrespective of the initial position of nucleosome *Y*, one obtains the value of *α* as 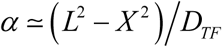 for *Y* < *X* and 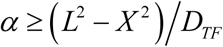 for *Y* > *X* and, *β* < 1 irrespective of the values of *Y*. One can compute the 1D diffusion coefficient as 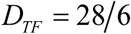 for *k* = 3 in the present context. These results are similar to the situation where 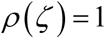 and 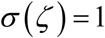 irrespective of the values of *ζ* as shown in **Fig. 4**.

To understand the effects of various types of potential functions on the 1D sliding mediated positioning-repositioning dynamics of nucleosomes in the absence of TFs, the integrals associated with *T*(*Y*) in **Eqs. 27** and **28** were numerically evaluated. For numerical integration we used the potential function 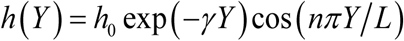 with *L* = 1000 bps and *h_0_* ~ 1 *k*_*B*_*T*. Further *n* was iterated from 0 to 5 and *γ* was iterated in *γ* ~ (0, 0.002, 0.004, 0.006, 0.008 0.01) bps^−1^. The generated stationary state nucleosome positioning profiles (**Figs. 7A** and **C**) and the respective MFPT values (**Figs. 7B** and **D**) are shown in **Figs. 7A-D**. Stationary state nucleosome positioning profiles were generated by imposing reflecting boundary conditions at *Y* = 0 as well as *Y* = 1000. For the purpose of MFPT computation we had set *Y* = 0 as a reflecting boundary and *Y* = 1000 as an absorbing boundary for the dynamics of nucleosome starting from *Y_0_*. Here number of peaks in the nucleosome positioning profiles seems to be approximately *n*/2. The MFPT decreases as *n* increases to 1 and 2 from *n* = 0. Thereafter MFPT steadily increases with *n*. These results clearly suggested that the overall MFPT associated with the sliding and crossing of a boundary by a nucleosome is positively correlated with the number of peaks in the occupancy profile as well as *h_0_* and negatively correlated with the exponent *γ* (**Fig. 7D**). These results are all logical since the extent of hindrance exerted on the repositioning dynamics of the nucleosome of interest is directly proportional to (1) the number other nucleosome roadblocks on its 1D diffusion path and (2) the free energy barrier associated with the transition of nucleosome from one minimum to another adjacent one which is directly proportional to the amplitude *h_0_*.

**FIG. 7.**
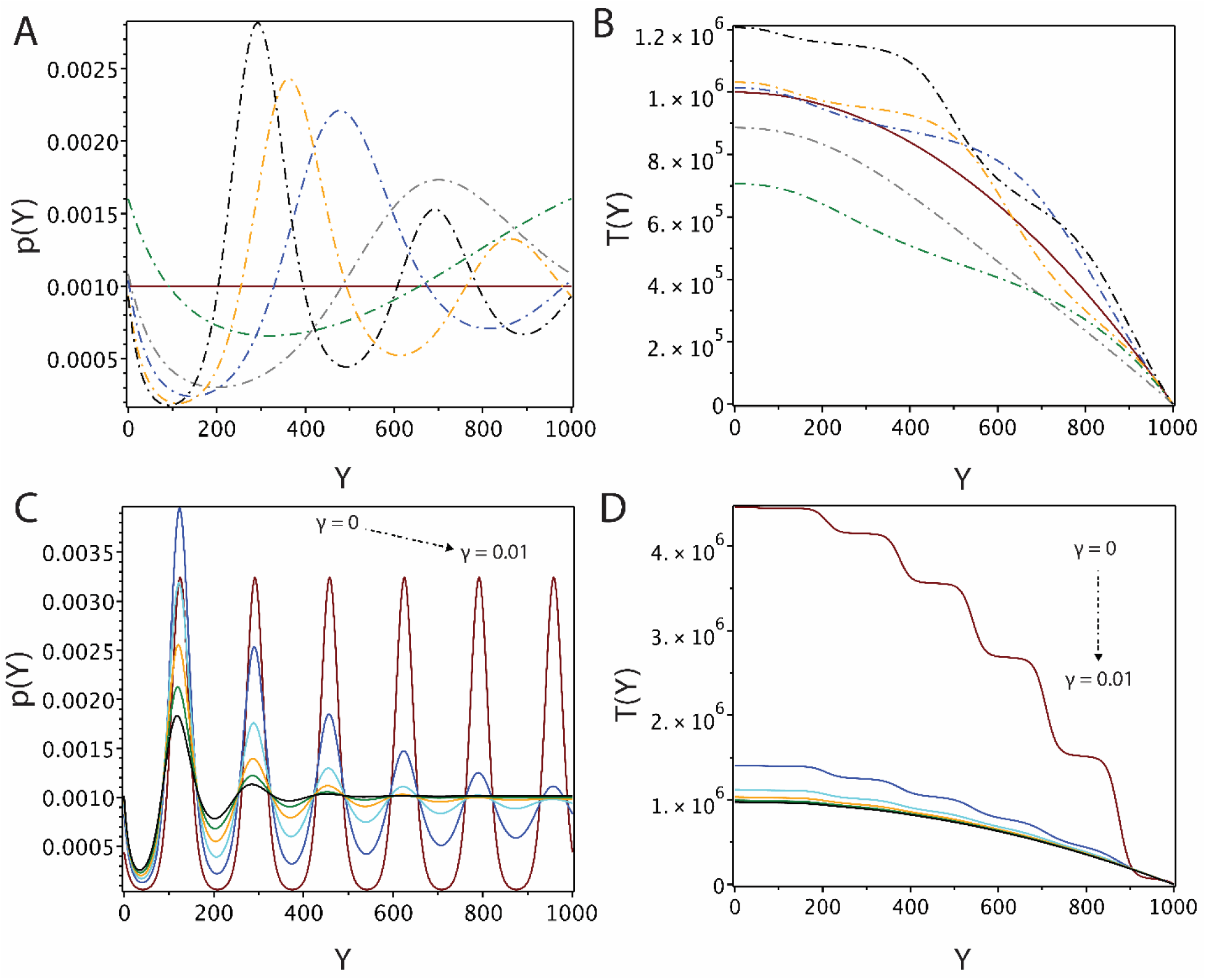
Effects of position dependent potential function 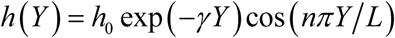on the positioning and sliding dynamics of nucleosomes along DNA in the absence of TFs. Here *Y* is measured in terms of bps and *T*(*Y*) is measured in terms of number of simulation steps. The MFPT *T*(*Y*) is the average time that is required by a nucleosome to slide through a DNA of length *L* bps. **Eqs. 27** (integrals in the expression for *T*(*Y*)) and **28** were numerically evaluated with *γ* ~ 0.002 bps^−1^, *L*= 1000 bps, *h_0_* ~ 1 *k*_*B*_*T* and *n* was iterated from 0 to 5. For the purpose of MFPT computation we set *Y* = 0 as a reflecting boundary and *Y* = 1000 as an absorbing boundary. Here the solid lines corresponds to *n* = 0. The MFPT decreases as *n* increases to 1 and 2 from *n* = 0 and thereafter MFPT steadily increases with *n*. A and B share common color i.e. MFPT denoted by green dotted line of B corresponds to occupancy profile denoted by green dotted line of A and so on. **A**. Stationary state nucleosome positioning profiles. **B**. Computed MFPT (*T*(*Y*)) values corresponding to each occupancy profile in **A**. Settings for **C** and **D** are *n* = 12, *L* = 1000, *h0* = 1 and *γ* was iterated through 0, 0.002, 0.004, 0.006 0.008 and 0.01 bps^−1^. **C**. Computed stationary state nucleosome positioning profiles. (**D)**. Computed MFPT (*T*(*Y*)) values associated with the escape of nucleosome from (0, *L*) corresponding to each occupancy profile in **C**.

## 4. Discussion

The 1D diffusion of TFs and ATP-independent sliding dynamics of nucleosomes are thermally driven stochastic processes. Here 1D sliding of nucleosomes along DNA is mainly achieved by the formation of thermal induced 10-bps bulge/loop defects or 1-bps twist defects in the DNA-nucleosome interaction network and their propagation via reptation dynamics of the DNA polymer [36, 39]. Predominantly, 1D sliding dynamics of nucleosomes seems to be driven by the 1-bps twist defect which induces a corkscrew type dynamics of nucleosomes within the wrapping of DNA [36, 37, 39]. On the other hand, the electrostatic attractive potential combined with the shielding effects of solvent ions creates a fluidic type environment at the DNA-TF interface which helps TFs to smoothly slide along DNA [7-9]. In the absence of other roadblocks and position dependent free energy potential profiles, the single bps movement of TFs and nucleosomes along DNA seems to be within microsecond and millisecond to second timescales respectively. The slow diffusion of nucleosomes along DNA is mainly due to the underlying wrapping-unwrapping dynamics of the DNA polymer which incurs huge free energy barrier. Our theoretical and simulation results put forward the following key observations on the collective dynamics of TF and nucleosome on the same DNA polymer.

(1) The roadblock effects of nucleosome are strongly dependent on the relative position of nucleosome with respect to TSS or TFBS. Especially the nucleosome exerts maximum amount of hindrance for the sliding dynamics of TFs when it is present in between TFs and their CRMs. This could be one of the reasons behind the occurrence of nucleosome depleted regions around TSSs and TFBSs [16, 52] under *in vivo* conditions. Otherwise organisms need to invest huge amount free energy input in the form of ATP hydrolysis to actively reposition the nucleosomes from the locations of TSS and CRMs. However the counter argument to this deduction can be as follows. The nucleosome depleted regions around TSSs or TFBSs are mainly due to the passive competition between TFs and nucleosome for binding with the same segment of DNA.

(2) When TFs land on the region of DNA that is located in between nucleosome and TSS or TFBS, then the search dynamics of TFs towards their CRMs will be enhanced to certain extent. However the extent of this enhancement effects is much lower than the extent of retardation effects of nucleosome when it is present in between TFs and their CRMs [51]. However TFs can make use of this enhancement effects of nucleosome only when the nucleosome depleted regions around CRMs are much longer than the typical sliding lengths of TFs which is around 100 bps in prokaryotic systems [11, 20, 58].

(3) TFs exhibit typical sub-diffusive type dynamics when they move close to the slowly diffusing nucleosomes. Such phenomenon allows TFs to stay close to nucleosomes for prolonged timescales which in turn is required for an efficient competitive binding of TFs and nucleosomes with the same stretch of DNA. However such sub-diffusive dynamics of TFs disappears as the distance between TFs and nucleosomes increases or the DNA is under condensed conformational state which in turn allows TFs to jump across nucleosomes.

(4) The roadblock effects of nucleosomes on the 1D diffusion dynamics of TFs can be almost nullified by the supercoiled or condensed conformational state of DNA which in turn favors hopping and intersegmental transfers. However TFs can completely overcome the retarding effect of nucleosomes only when the degree of condensation of DNA is such that TFs can easily hop over the nucleosome roadblocks transfers i.e. TFs require a minimum jump size of *k* > 147 bps.

Contrasting from the occupancy pattern of nucleosomes around TFBS, the occupancy pattern of TFs around their corresponding CRMs shows a maximum at the location of CRM which declines as the distance of TF from the location of its CRM increases. Such occupancy pattern of TFs around their CRMs seems to be achieved in two possible ways viz. (1) by manipulating the relative distribution of the sequence mediated traps around the CRMs [17] and (2) by attenuating the 1D diffusion coefficient along DNA as the TF molecule approaches its CRM which is mainly achieved by fine tuning the conformational fluctuations at the DBDs of TFs [20]. Detailed computational studies suggested that the sequence mediated traps around the location of CRMs are positioned [17] in the natural systems such that relatively strong traps are positioned close to the location of CRMs and weak traps are positioned throughout the genomic DNA. This in turn decreases the effective 1D diffusion coefficient as the TF molecule moves close to its CRM.

Earlier studies suggested that DBDs of TFs fluctuate between at least two different conformational states viz. stationary and mobile which are characterized by distinct 1D diffusion coefficients [19]. For the purpose of convenience we denote them here as *χ*_*TF,S*_ and *χ*_*TF,M*_ respectively for stationary and mobile conformational states. The free energy barrier associated with the fluctuations among these two conformational states seems to be <3*k*_*B*_*T* [53]. This resembles the dynamics of downhill folding proteins [54] at their mid-point denaturation temperatures. The stationary conformational state of DBDs is more sensitive to the sequence information of DNA than the mobile state but they diffuse slowly. The mobile conformational state of DBDs of TFs is less sensitive to the sequence information but they diffuse rapidly along the DNA lattice. Clearly we have *χ*_*TF,S*_ ≤ *χ*_*TF,M*_ by definition. When TFs move close to their CRMs then the extent of conformational fluctuations in their DBDs decreases in a monotonic manner and subsequently the DBDs of TFs form a tight complex with lowest possible conformational fluctuations upon finding their CRMs [19]. Such decrease in the extent of conformational fluctuations at the DBDs may also be linked with the presence of strong sequence traps as TFs move close towards their CRMs [17].

Recent theoretical studies [20] suggested that when the rate of flipping between the stationary and mobile conformational states of DBDs increases towards infinity, then the overall effective 1D diffusion coefficient of TFs transform from *χ*_*TF,G*_ = 2*χ*_*TF,S*_ *χ*_*TF,M*_ / (*χ*_*TF,S*_ + *χ*_*TF,M*_) to *χ*_*TF,A*_ = (*χ*_*TF,S*_ + *χ*_*TF,M*_) / 2. Here *χ*_*TF,A*_ is the arithmetic mean of the 1D diffusion coefficients *χ*_*TF,S*_ and *χ*_*TF,M*_, and *χ*_*TF,G*_ is their geometric mean. Clearly we have *χ*_*TF,G*_ ≤ *χ*_*TF,A*_. That is to say the 1D diffusion coefficient of TFs decreases as TFs move close to their cognate sites. Contrasting from the dynamics of DBDs of TFs, the extent of breathing dynamics of nucleosome particles increases as they move close to CRMs which leads to rapid diffusion of nucleosomes over CRMs. That is to say, natural systems are designed such that the dynamics of TFs becomes slow as they move close to their CRMs but the dynamics of nucleosomes becomes rapid as they move across those CRMs. As a result of such sequence dependent fluctuations in the DBDs of TFs as well as breathing dynamics nucleosomes, one finally observes a maximum occupancy of TFs and minimum occupancy of nucleosomes at the CRMs of TFs throughout the natural systems.

The key assumptions in the derivation of **Eqs. 17-19** are (1) both TF and nucleosome perform a normal diffusion along DNA and (2) nucleosome which crossed the location of CRM (*Y* = *L*) will not return back into the interval (0, *L*) and impedes the dynamics of TF further. Experiments revealed the sub-diffusive type dynamics of nucleosomes under crowded environments [45, 46]. Our stochastic simulations suggested that the sub-diffusive type dynamics of TFs occurs only when TFs move close to nucleosomes. It seems that such anomalous type diffusion disappears as the distance between TF and nucleosome increases or the DNA template of interest is under condensed conformational state which can allow a jump size *k* ≥ 150 bps [57]. Since the average length of the linker DNA that connects two consecutive nucleosomes is in the range ~10-100 bps [28, 30], one can safely ignore the hindrance effects on MFPT due to sub-diffusive dynamics of TFs as they move close to nucleosomes.

## 5. Conclusion

Site-specific binding of transcription factors with their *cis*-regulatory modules (CRMs) located on the genomic DNA is critical for the precise regulation of eukaryotic genes. It is an established fact that TFs locate their cognate sites of DNA via a combination of 1D and 3D diffusion routes. The overall search time required by TFs to locate their targets under *in vivo* conditions is strongly influenced by the factors viz. conformational state of DNA, fluctuations in the DBDs of TFs, electrostatic forces present at the DNA-protein interface, random occurrence of sequence traps and, semi-stationary roadblocks such as nucleosomes. Nucleosomes play several critical roles ranging from packaging of the genomic DNA to regulation of various genes. Unlike prokaryotic systems, TFs in eukaryotes need to search for their cognate sites in the presence of slowly sliding nucleosome roadblocks on the same DNA.

Using random walk models, here we have shown that nucleosomes can efficiently control the relative search times spent by TFs on 1D and 3D diffusion routes towards finding their cognate sites on DNA. The roadblock effects of nucleosomes seems to be dependent on the relative position of nucleosome with respect to TFs and their CRMs. Nucleosomes exert maximum amount of hindrance to the 1D diffusion dynamics of TFs when they present in between TFs and their cognate sites. The effective 1D diffusion coefficient (*χ*_*TF*_) associated with the dynamics of TFs in the presence of nucleosomes seems to decrease with the free energy barrier (*µ*) associated the sliding dynamics of nucleosomes as 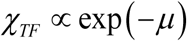. As a result the average time (*η*_*L*_) that is required by TFs to scan *L* number of binding sites on DNA via 1D diffusion increases with *μ* as 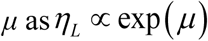. When TFs move close to nucleosomes, then they exhibit a typical sub-diffusive type dynamics which disappears as the distance between TFs and nucleosomes increases.

Nucleosomes can enhance the search dynamics of TFs when TFs present in between nucleosomes and transcription factor binding sites (TFBS). The level of such enhancement effects seems to be much lesser than the level of retardation effects when nucleosomes presence in between TFs and their cognate sites. These results suggested that nucleosome depleted regions around the cognate sites of TFs is mandatory for an efficient site-specific interactions of TFs with DNA. In line with our predictions, the genome wide positioning pattern of TFs shows maximum at their specific binding sites and the positioning pattern of nucleosome shows minimum at those locations under *in vivo* conditions. We argued that this could be a consequence of increasing level of breathing dynamics of nucleosomes and decreasing levels of fluctuations in the DNA binding domains of TFs as they move across TFBS. Here the extent of breathing dynamics of nucleosomes and fluctuations in the DBDs of TFs are positively correlated with their respective 1D diffusion coefficients. As a result, the dynamics of TFs becomes slow as they approach their cognate sites so that TFs form tight site-specific complex and the dynamics of nucleosomes becomes rapid so that they rapidly pass through the cognate sites of TFs. Several *in vivo* data on the positioning pattern of nucleosomes as well as TFs along the genomic DNA seem to agree well with our arguments. The condensed conformational state of DNA significantly decreases the retarding effects of nucleosome roadblocks. The retarding effects of nucleosomes on the 1D diffusion dynamics of TFs can be nullified when the degree of condensation of DNA is such that the jump size associated with the dynamics of TFs is *k* > 150 bps.

